# Coxsackievirus B infection invokes unique cell-type specific responses in primary human pancreatic islets

**DOI:** 10.1101/2024.07.23.604861

**Authors:** Daniel A. Veronese-Paniagua, Diana C. Hernandez-Rincon, Jared P. Taylor, Hubert M. Tse, Jeffrey R. Millman

## Abstract

Coxsackievirus B (CVB) infection has long been considered an environmental factor precipitating Type 1 diabetes (T1D), an autoimmune disease marked by loss of insulin-producing β cells within pancreatic islets. Previous studies have shown CVB infection negatively impacts islet function and viability but do not report on how virus infection individually affects the multiple cell types present in human primary islets. Therefore, we hypothesized that the various islet cell populations have unique transcriptional responses to CVB infection. Here, we performed single-cell RNA sequencing on human cadaveric islets treated with either CVB or poly(I:C), a viral mimic, for 24 and 48 hours. Our global analysis reveals CVB differentially induces dynamic transcriptional changes associated with multiple cell processes and functions over time whereas poly(I:C) promotes an immune response that progressively increases with treatment duration. At the single-cell resolution, we find CVB infects all islet cell types at similar rates yet induces unique cell-type specific transcriptional responses with β, α, and ductal cells having the strongest response. Sequencing and functional data suggest that CVB negatively impacts mitochondrial respiration and morphology in distinct ways in β and α cells, while also promoting the generation of reactive oxygen species. We also observe an increase in the expression of the long-noncoding RNA *MIR7-3HG* in β cells with high viral titers and reveal its knockdown reduces gene expression of viral proteins as well as apoptosis in stem cell-derived islets. Together, these findings demonstrate a cell-specific transcriptional, temporal, and functional response to CVB infection and provide new insights into the relationship between CVB infection and T1D.

## Introduction

Type 1 diabetes (T1D) is a chronic condition marked by high blood glucose levels caused by an autoimmune response that targets and kills insulin-secreting pancreatic β cells. This disease currently affects roughly 27-54 million people worldwide.^1^ Predisposition to T1D is caused by genetic susceptibility in multiple loci, including *INS, IFIH1*, and the human leukocyte antigen (HLA) genes.^2^ Yet, genetic background alone does not explain neither the 35% discordant rate for T1D among monozygotic twins nor the doubling of the annual incidence rate in the last twenty years to an estimated 3.9%.^2–4^ Multiple groups have turned to environmental factors, such as microbial diversity, diet, and stressors, as triggers for β cell autoimmunity and dysfunction.^4–13^ However, our incomplete understanding of the relationship between the complex initiation and progression of T1D and the environment have led to a deficit of proper therapeutics to protect β cells from the immune system. Therefore, there is a great demand to investigate these environmental triggers with improved human *in vitro* models to further elucidate this relationship and ultimately discern methods to delay or prevent β cell autoimmunity.

For decades, enterovirus infection has been stipulated as an environmental trigger associated with T1D.^2,14^ These nonenveloped viruses, which belong to the *Picornaviridae* family, possess a single-stranded RNA genome of approximately 7.5 kilobases that encodes 11 proteins. This family encompasses various pathogens, including coxsackieviruses, polioviruses, and rhinoviruses.^15^ Enteroviral capsid proteins have previously been detected in 60-70% of pancreata and islets of Langerhans from T1D patients compared to 6% in non-diabetic control groups.^2,4–6,15–17^ A prior study revealed that coxsackieviruses accounted for 64% of the enteroviruses identified in excess in the stool of T1D susceptible patients, which were detectable up to a year prior to the first detection of islet autoantibodies.^18^ Coxsackieviruses are organized into 23 group A viruses, which are mostly tropic for striated muscle in mice, and six group B viruses, which are tropic for the pancreas, heart, liver, brown fat, central nervous system, and striated muscle.^19^ More specifically, group B coxsackievirus (CVB) strains have been shown to cause persistent, chronic inflammatory infections that can lead to pancreatitis, myocarditis, and meningitis in both humans and mice.^20–22^

While many groups have reported on the association of CVB with T1D, there is limited information on how CVB infection impacts the various cell types in human pancreatic models. Using rodent and human β cell lines as well as primary human islets, previous reports have revealed that multiple CVB variants induce endoplasmic reticulum and Golgi stress, cell degeneration, pyknosis, reduction of islet identity genes, and decreased insulin content and secretion.^21,23–26^ CVB4- and CVB5-induced type 1 interferon (IFN) signaling was shown to be stronger in rat α cells compared to β cells while persistent CVB1 infection of human PANC-1 ductal cells induced differential expression of genes associated with immune response, extracellular matrix, β cell-to-cell communication, and hormone secretion.^26,27^ However, a global transcriptional response and cell-type specific responses have not been investigated in CVB-infected human primary islets.

Moreover, CVB tropism at a cellular level in humans has been primarily elucidated through infection of cell lines, analysis of entry receptor expression, and the colocalization of viral and islet cell identity protein markers. However, these methods are limited considering CVB infects non-receptor cells through exosomes in non-pancreatic tissues.^28^ Studying such interactions in mouse models is difficult since CVB only infects exocrine cells in the murine pancreas.^19^ Also, obtaining data beyond the effects of CVB infection at the whole islet level has been limited by the heterogenous composition of islets and the inability to distinguish responses of specific cell types. Therefore, whether CVB infects and persists in all human pancreatic endocrine tissue, differentially affects islet cell types, or triggers islet autoimmunity/T1D remains undetermined.

In this study, we investigate the impacts of CVB infection on a 3-dimensional human *ex vivo* pancreatic islet organoid model. We treated whole islets with either CVB3 or poly(I:C) for 24 hr and 48 hr and performed single-cell RNA sequencing on these samples and their respective control. Through genetic cell marker expression, we identified pancreatic endocrine and exocrine cells. We compared whole islet response to the different stressors using differential gene expression analysis and identified on cell-type specific transcriptional signatures following CVB3 infection. We followed up our observations through assessments of mitochondrial morphology and function as well as characterization of a novel gene, *MIR7-3HG*, in islet response to infection. Our work elucidates unique endocrine and exocrine cell-type specific responses to CVB3 infection and highlights the importance of investigating viral infection at a tissue level to develop novel therapies that alleviate damage from such pathogens.

## Results

### Poly(I:C) and CVB3 both induce immune responses in islets yet differ in magnitude of transcriptional response and temporal effects

We set out to investigate CVB infection of human pancreatic islets to identify any impacts on transcriptional signature at a single-cell resolution. To do this, we performed independent time course experiments treating primary human islets from four cadaveric donors for 24 and 48 hours with either water, CVB3-eGFP at MOI 20, or polyinosinic-polycytidylic acid (poly(I:C))—a synthetic double-stranded RNA mimicking viral infection (Extended Data Fig. 1a, Supplementary Data 1). After treatment, these islets were collected and processed for single-cell RNA sequencing (scRNA-seq) to obtain the transcriptional signature of the various islet cell types (Fig. 1a). We also collected RNA from three of the four cadaveric islets and verified CVB3 infection by measuring viral RNA presence via real-time quantitative PCR (rt-qPCR) (Fig. 1b, Supplementary Data 1). Following quality check and filtering, we analyzed 31,446 cells across all six conditions (Extended Data Fig. 1b). Gene marker expression indicated the presence of all major endocrine islet cell types as well as other cells common to the pancreatic microenvironment, such as ductal, acinar, and endothelial cells, in all conditions (Fig. 1c-d, Extended Data Fig. 1c-f).

**Fig. 1.**
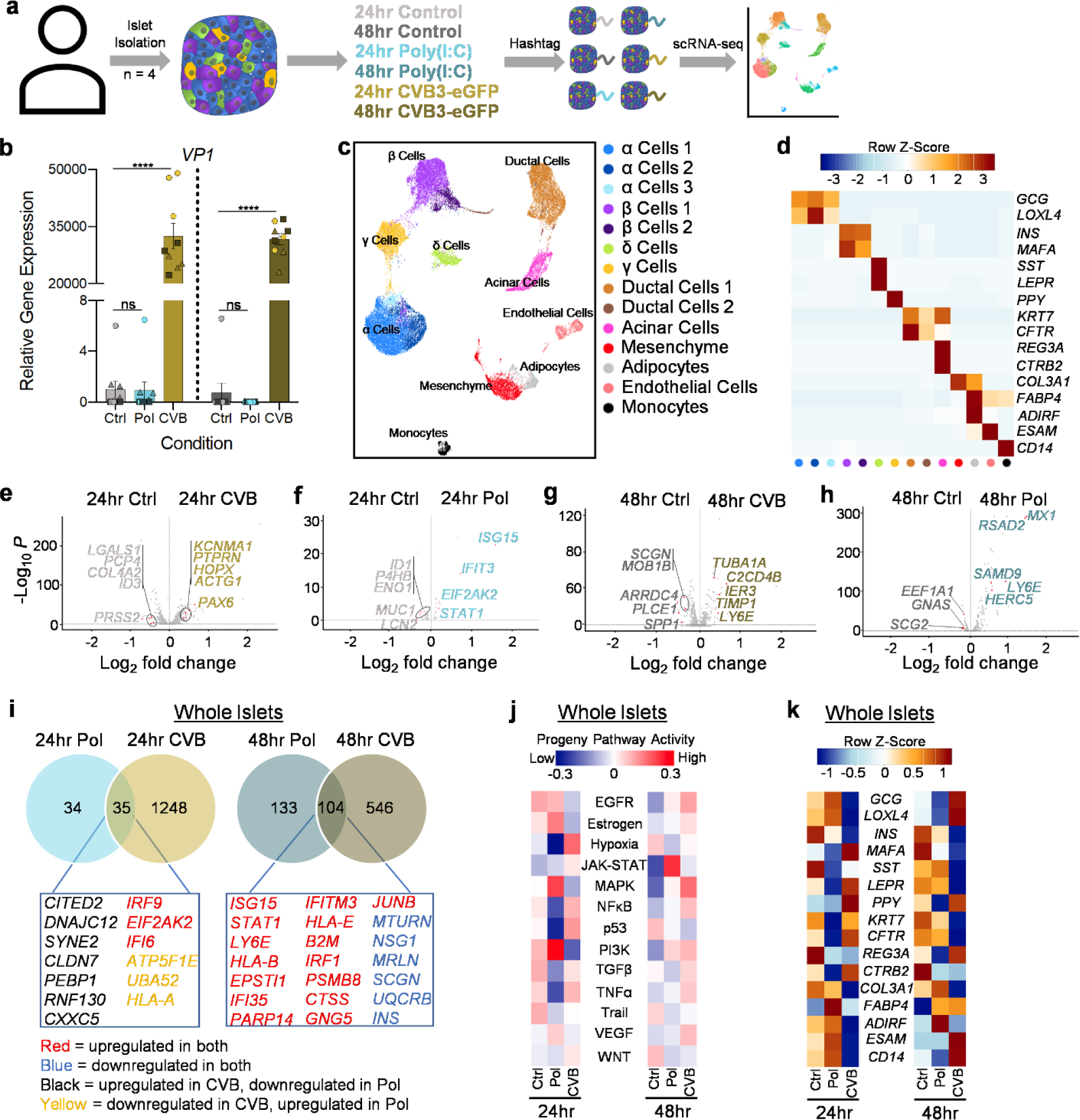
Single-cell RNA sequencing of primary human islets reveals unique transcriptional responses to CVB3 and poly(I:C) treatment. **a**, Schematic depicting workflow to generate scRNA-seq data from islets isolated from cadaveric donors (n=4). **b**, rt-qPCR of viral antigen gene expression to show infection of whole islets (n = 9. Patient 01 = Triangle, Patient 02 = Square, Patient 03 = Circle). Error bars represent the s.e.m. **c**, UMAP of different cell populations identified upon integration of scRNA-seq data from all patients. **d**, Heatmap depicting genetic marker expression for each cluster. **e-h**, Volcano plots highlighting the differences in whole islet gene expression when comparing (**e**) Control to CVB3 24 hr (total variables = 1284), (**f**) Control to poly(I:C) 24 hr (total variables = 69), (**g**) Control to CVB3 48 hr (total variables = 651), and (**h**) Control to poly(I:C) 48 hr (total variables = 237). **i**, Venn diagrams comparing statistically significant DEG in whole islets treated with either poly(I:C) or CVB3 at 24 hr and 48 hr. DEG for each condition was obtained comparing the treatment to its respective control. **j**, Progeny analysis of 14 major pathways showing the scaled pathway activity scores across conditions over time in primary human islets. Negative values correspond to decreased pathway activity and positive values correspond to increased pathway activity. **k**, Heatmap depicting differences in marker expression across conditions over time in primary human islets. *** = Ordinary One-way ANOVA with Tukey’s multiple comparisons.

We performed pairwise differential gene expression analysis by comparing each treatment to its respective control groups to identify genes associated with viral response. At 24 hr, CVB3 and control groups variably expressed 1,284 genes (Fig 1e, Supplementary Data 2). Within the analysis, *PTPRN, PAX6, KCNMA1, HOPX*, and *ACTG1* were enriched in CVB3-infected islets whereas genes enriched in control cells included *LGALS1, PCP4, COL4A2, ID3*, and *PRSS2* (Fig. 1e). Many of these genes are involved in various cell processes, including calcium signaling, transcription regulation, and protein trafficking. Gene ontology (GO) analyses for genes enriched in CVB3-infected cells revealed associations with insulin secretion regulation, vesicle-mediated transport, transcription, protein transport, apoptosis, and stress response whereas control-enriched genes were associated with integrin-mediated signaling, cellular respiration, endocytosis, and the extracellular matrix (ECM) (Extended Data Fig 3a-b, Supplementary Data 3).

At 24 hr, poly(I:C) and its control differed in 69 genes (Fig 1f, Supplementary Data 2). Poly(I:C) treatment induced expression of genes associated with a double-stranded RNA-mediated immune response, including *ISG15, IFIT3, STAT1*, and *EIF2AK2* (Fig. 1f). In contrast, control cells exhibited higher expression of *ID1, P4HB, ENO1*, and *LCN2*, which are involved in transcription, post-translational processes, glycolysis, and lipid transport, respectively (Fig. 1f). *MUC1* was also present in control cells (Fig. 1f). The GO terms for genes upregulated in poly(I:C)-treated cells were associated with translation, immune response, and endothelial cell migration (Extended Data Fig 3c, Supplementary Data 3). We did not observe GO terms with a significant adjusted p-value (<0.05) for genes downregulated in 24 hr poly(I:C) conditions. 651 variable genes were detected when comparing CVB3 and control groups at 48 hr (Fig. 1g, Supplementary Data 2). Relative to CVB3 infection, control cells had higher expression of *SCGN, MOB1B, ARRDC4, PLCE1*, and *SPP1*, which are genes associated with calcium signaling, negative regulation of cell proliferation, protein transport, PI3K signaling, and cell-matrix interaction (Fig. 1g). GO analyses on control-enriched genes showed associations with translation and cellular respiration (Extended Data Fig. 4b, Supplementary Data 3). In contrast, CVB3 infected cells overexpressed *TUBA1A, C2CD4B, IER3, TIMP1*, and *LY6E*, which play roles in the cytoskeleton, antiviral response and apoptosis, the ECM, and viral entry (Fig. 1g). GO analyses on CVB3-enriched genes highlighted terms associated with inflammatory response, apoptosis, proliferation, cellular respiration, and ubiquitin ligases (Extended Data Fig. 4a, Supplementary Data 3).

When comparing control cells to poly(I:C)-treated cells at 48 hr, we observed 237 variable genes (Fig. 1h, Supplementary Data 2). *EEF1A1, GNAS*, and *SCG2* were highly expressed in control cells. These genes are involved in translation, GPCR signaling, and hormone secretion (Fig. 1h). Poly(I:C)-treated cells expressed antiviral genes, such as *MX1, RSAD2, SAMD9, LY6E*, and *HERC5* (Fig. 1h). GO analyses of genes upregulated with poly(I:C) treatment also generated terms associated with antiviral response and translation (Extended Data Fig. 4c-d, Supplementary Data 3).

Given the substantial variation in gene expression when comparing control cells to either CVB3- or poly(I:C)-treatment, we delineated genes that are shared and those that are unique to the two treatment groups. To achieve this, we first took the gene lists generated from pairwise comparisons with control conditions at each timepoint and removed *eGFP*. Treatment-enriched genes were classified as upregulated while control-enriched genes as downregulated in treated cells. To identify shared genes between the lists, we generated Venn diagrams, which suggest poly(I:C) and CVB3-treated cells share 35 genes at 24 hr and 104 genes at 48 hr (Fig. 1i, Supplementary Data 4). At 24 hr, there are several shared genes upregulated in CVB3 but downregulated in poly(I:C)— such as *CITED2, PEBP1*, and *CXXC5*—that regulate hypoxia, ubiquitination as well as MAPK, NF-κB, and Wnt signaling (Fig. 1i, Supplementary Data 4). On the other hand, *ATP5F1E* and *UBA52*, which are involved in mitochondrial function and ubiquitination, are downregulated in CVB3 but upregulated in poly(I:C).

In cells treated for 48 hr, *MTURN, NSG1, MRLN, SCGN*, and *UQCRB* were downregulated in both conditions (Fig. 1i, Supplementary Data 4). These genes regulate multiple processes including NF-κB, MAPK, apoptosis, calcium flux in the endoplasmic reticulum (ER), and respiration. In both treatment conditions and timepoints, we found upregulation of several antiviral and immune response genes, including *IRF9, IFI6, EPSTI1*, and *CTSS*.

To highlight the differences between CVB3- and poly(I:C)-treated cells, we performed pairwise comparisons of the treatment groups at each timepoint. For the 24 hr timepoint, our analysis revealed 2,753 variable genes (Extended Data Fig. 2a, Supplementary Data 2). In CVB3-infected cells, we identified genes associated with calcium signaling (*TACSTD2*), exosome biogenesis (*SDC4*), and actin polymerization (*TMOD1*) and found GO terms related to protein transport and degradation, vesicle-mediated transport, autophagy, ER stress, the cytoskeleton, and mTORC1 signaling (Extended Data Fig. 2a, Extended Data Fig. 5a, Supplementary Data 2, Supplementary Data 3). *SPARC, PRSS2*, and *TIMP1*, which are involved in ECM degradation, were all enriched in poly(I:C)-treated cells (Extended Data Fig. 2a). GO analysis highlighted genes associated with cellular respiration, ECM, translation, and lysosomes (Extended Data Fig. 5c, Supplementary Data 3).

At 48 hr post treatment, there were 438 variable genes when comparing CVB3- and poly(I:C)-treated cells to each other (Extended Data Fig. 2b, Supplementary Data 2). CVB3-infected cells exhibited higher expression of multiple inflammatory response genes—*CEBPD, TMEM176B*, and *C2CD4B*—as well as *SOD2* and *PTP4A3*, which regulate reactive oxygen species (ROS) production and cell proliferation, respectively (Extended Data Fig. 2b). The GO terms associated with genes enriched in the CVB3 condition included translation and mitochondrial stress (Extended Data Fig. 5b, Supplementary Data 3). In poly(I:C)-treated cells, there was enrichment of antiviral response genes, such as *MX1, RSAD2*, and *STAT1*, and regulators of proliferation— *EGR1* and *FOS*. GO analyses of enriched genes with poly(I:C) treatment indicated associations with gene expression, lysosomal, and viral response programs (Extended Data Fig. 2b, Extended Data Fig. 5d, Supplementary Data 3).

Next, we sought to investigate how islets were impacted over time by each treatment. To do this, we leveraged the progeny package to better understand the effects of each treatment on major signaling pathways.^29^ Progeny takes the directionality of gene expression to compute scaled pathway activity scores across conditions. Our analysis highlighted differential impacts on cell signaling when comparing control, poly(I:C), and CVB3. For instance, hypoxia activity was increased in CVB3 and decreased in poly(I:C) when compared to control at 24 hr. By 48 hr, these scores were alleviated (Fig. 1j, Supplementary Data 5). We also saw temporal differences in PI3K, Estrogen, and MAPK signaling. CVB3 infection induced higher activity scores for TNFα and NF-κB signaling at both timepoints while JAK-STAT was drastically higher in poly(I:C) conditions (Fig. 1j, Supplementary Data 5). We observed higher average expression of genes involved in type 1 IFN signaling in poly(I:C)-treated cells that is increased by 48 hr (Extended Data Fig. 2c).

When comparing poly(I:C) and CVB3 treatment in these cell types, we noticed they impacted expression of cell identity markers differently over time (Fig. 1k). For instance, α cell identity was increased by 24 hr in poly(I:C) but then decreased at 48 hr. In contrast, α cell identity was negatively impacted by CVB3 at 24 hr but saw improvement by 48 hr. Mesenchymal identity marker expression was decreased at both timepoints for poly(I:C) and CVB3. In β cells, *INS* and *MAFA* expression was decreased at both timepoints with poly(I:C) treatment. Although *INS* was downregulated in CVB3-treated cells as well, we report *MAFA* expression was increased at 24 hr but was decreased by 48 hr. A similar trend with CVB3 treatment was observed with δ cell markers *SST* and *LEPR* while expression of both markers decreased at first and then increased by 48 hr with poly(I:C) (Fig. 1k). Acinar cell identity was negatively impacted by poly(I:C) whereas CVB3 treatment decreased *REG3A* expression at 24 hr, but its expression was increased after 48 hr. We do not see a similar trend for *CTRB2*, another acinar cell marker, in CVB3-infected cells. Finally, adipocyte identity increased at 24 hr and 48 hr with poly(I:C) treatment while we saw decreases with CVB3 at 24 hr with an increase in *FABP4* expression after 48 hr.

### CVB3 infects all islet cell types yet induces unique cell-type specific transcriptional responses

We set out to determine if CVB3 infects all the various islet cell types. We subsetted the CVB3 conditions and independently measured which cell types were eGFP^+^ and at what frequency. We found eGFP expression in all the cell types in our dataset (Extended Data Fig. 6a). We calculated 62.5% to 87.1% of cells were eGFP^+^ at 24 hr while 69.7% to 89.8% were eGFP^+^ at 48 hr (Fig. 2a, Supplementary Data 6). α, γ, and ductal cells were among the highest rated eGFP^+^ cells. We next measured the mean gene expression of the CVB entry receptors, *CD55* and *CXADR*. All cell types except acinar, mesenchyme, adipocytes, and monocytes expressed *CD55* at moderate levels (Fig. 2b). However, only δ, ductal, and acinar cells highly expressed *CXADR* (Fig. 2c). Thus, our data indicate only δ and ductal cells express both *CD55* and *CXADR*.

**Fig. 2.**
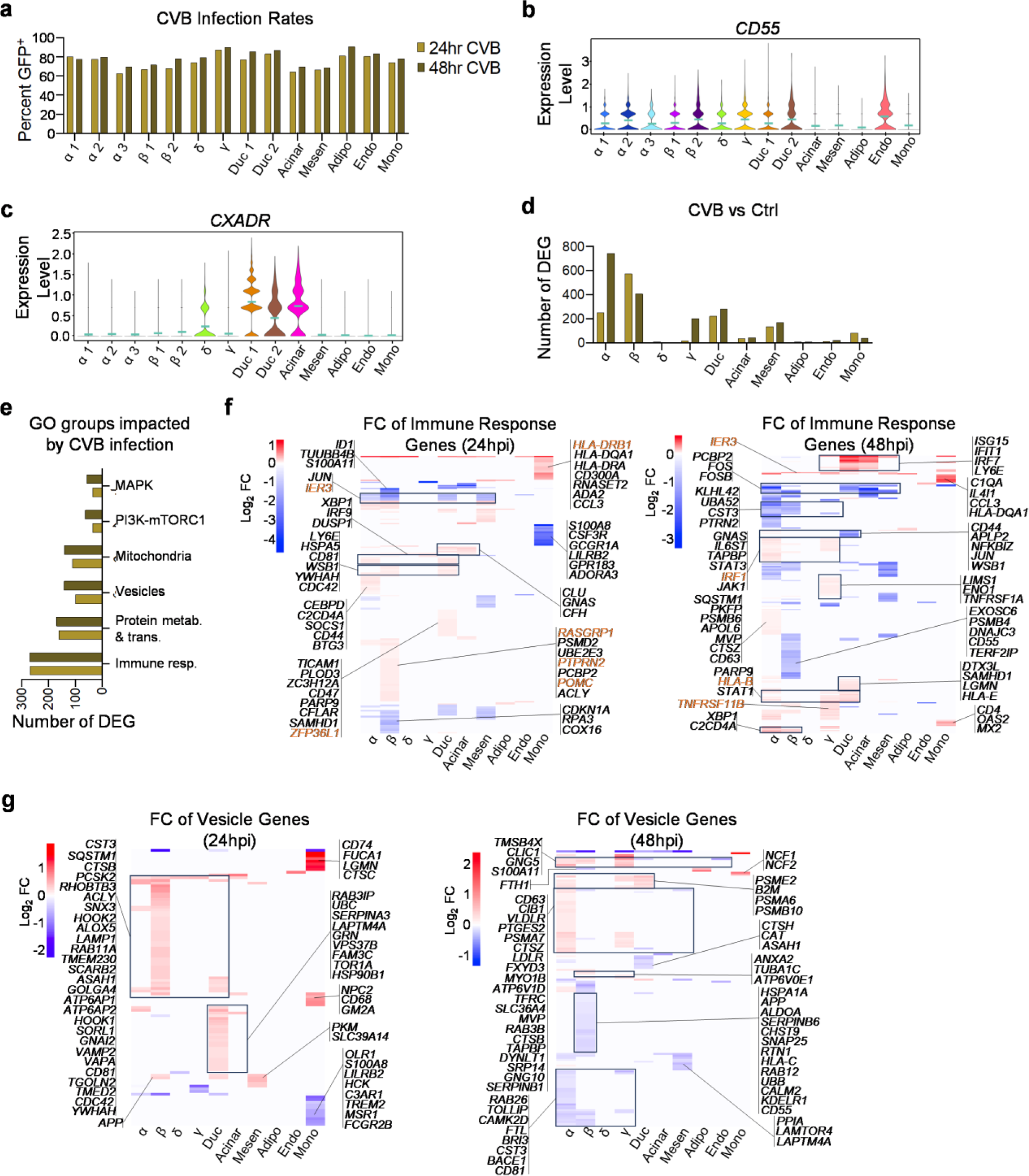
scRNA-seq reveals unique cell-type specific responses to CVB3 infection despite similar infection rates. **a**, Bar graph of CVB3 infection rates across cell types at 24 hr and 48 hr. **b, c**, Violin plots of CVB entry receptors (**b**) *CD55* and (**c**) *CXADR* across cell types in all treatment conditions. **d**, Bar graph showing the number of DEGs for each CVB3 timepoint when compared to their respective controls. **e**, Bar graph showing the number of DEG for selected themes of common Gene Ontology terms for each CVB3 timepoint compared to their respective controls. **f**, Heatmaps depicting log_2_ fold change (FC) of immune response-associated genes when comparing each CVB3 timepoint to their respective controls (orange font = T1D associated genes). **g**, Heatmaps depicting log_2_ fold change of vesicle-associated genes when comparing each CVB3 timepoint to their respective controls.

Next, we investigated how CVB3 infection differentially affects gene expression in the several islet cell types. First, we subsetted each cell type and performed pairwise comparisons using CVB3-infected cells and their corresponding control to identify differentially expressed genes (DEGs). α, β, and ductal cells had the highest number of DEGs with α and ductal cells displaying an increase between 24 hr and 48 hr while β cells had a decrease (Fig. 2d, Supplementary Data 7). On the other hand, δ cells, adipocytes, endothelial cells, and acinar cells were among the groups with the lowest number of DEGs. Despite these observations, we did not measure drastic differences in cell composition by 48 hr for each cell type except a small reduction in δ cells (Extended Data Fig. 6b, Supplementary Data 8).

We also explored the impact of CVB3 infection on cellular processes for each cell type. To achieve this, we first performed independent GO analyses for each cell type at both timepoints (Supplementary Data 9). We then categorized statistically significant GO terms into groups describing various processes and structures, such as gene expression, mitochondria, vesicles, and immune response. We dubbed these groups as GO themes (Supplementary Data 10). For each timepoint, we next listed the genes associated with the corresponding GO terms for the individual cell types and ran pairwise comparison between control and CVB3- infected cells to determine the number of DEGs with statistically significant fold changes (Fig. 2e, Extended Data Fig. 6c, Supplementary Data 11). The GO themes with the highest number of DEGs at 24 hr and 48 hr were associated with immune response [269 & 270 genes], gene expression [201 & 141 genes], protein metabolism and transport [162 & 171 genes], mitochondria [111 & 141 genes], and vesicles [100 & 142 genes] (Fig. 2e, Extended Data Fig. 6c).

We did not observe GO themes unique to specific cell types. Heatmaps for immune response-associated genes revealed gene expression unique to various cell types as well as shared genes (Fig. 2f, Supplementary Data 11). At both timepoints, the cells with the most differential transcriptional signatures were α, β, and ductal cells. At 24 hr, these three cell populations had mostly upregulated genes—sharing some including *JUN, WSB1, YWHAH*, and *CDC42*—and downregulated various ribosomal protein genes (Fig. 2f). We also observed β and ductal cells upregulated *LY6E* and *CD81*, which are associated with viral entry. Ductal and α cells also upregulated various type 1 IFN- and immune-associated genes including *IRF9, NFKBIZ*, and *IRF1* for both cell types, *TICAM1* and *PARP9* in ductal cells, and *CEBPD, C2CD4A, SOCS1*, and *STAT3* in α cells (Fig. 2f, Supplementary Data 11).

With 127 DEGs, β cells had the biggest immune response at 24 hr when comparing CVB3-infected cells to control. Infected β cells upregulated *RASGRP1, PTPRN2*, and *POMC*, which are all associated with T1D (Fig. 2f). On the other hand, most of the differentially regulated immune response-associated genes in infected β cells were downregulated at 48 hr including *CD81, HLA-A, CD55*, and *HLA-C* (Fig. 2f, Supplementary Data 11). *IER3*, which is associated with T1D, remained upregulated in CVB3-infected α, β, and mesenchymal cells compared to their 24 hr counterparts but was also upregulated in γ and δ cells by 48 hr. Many of the class II HLAs upregulated in monocytes at 24 hr were not differentially expressed at 48 hr (Fig. 2f, Supplementary Data 11).

Ductal and acinar cells had the strongest IFN-associated response upon CVB3 infection with shared upregulation of at least 14 genes, including *ISG15, IFIT1, IRF7, IFI35*, and *OAS1* (Fig. 2f, Supplementary Data 11). Additionally, ductal cells depicted increased expression of *HLA-B, PARP9, STAT1, IRF9*, which were also upregulated in α and γ cells, as well as *HLA-E, TAP1*, and *B2M* (Fig. 2f, Supplementary Data 11). By 48 hr, CVB3-infected α cells underwent an increase in DEGs compared to 24 hr with downregulation of *FOS, FOSB, DUSP1*, and *PTEN* and upregulation of *TAPBP, STAT3, HSPA5, XBP1, IRF1, JAK1*, and *HLA-E* (Fig. 2f, Supplementary Data 11). At 48 hr post infection, γ cells depicted similar trends to α cells with increased expression of translation-associated genes as well (Fig. 2f, Supplementary Data 11).

We noted at both timepoints that many of the differentially expressed α and β cell genes associated with the immune response theme encode ubiquitin components and products involved in protein degradation and the Unfolded Protein Response (UPR) with an overall decrease in expression of ribosomal-associated genes. As a result, we evaluated the GO themes associated with vesicles, lysosomes, and autophagy. At 24 hr, we observed β and ductal cells have the strongest induction of vesicle-associated genes across all islet cell types although there was little transcriptional overlap between them (Fig. 2g, Supplementary Data 11). In β cells, there was increased expression of several genes associated with vesicle as well as protein and small molecular trafficking, including *RHOBTB3, SNX3, HOOK2, TMEM230*, and *VAMP2*. β cells also increased expression of *SQSTM1, LAMP1, SCARB2, ATP6AP2*, and *WIPI1*, which are all associated with lysosomes and autophagy, as well as protease expression through *CTSB* and *PCSK2* (Fig. 2g, Supplementary Data 11). Both ductal and β cells upregulated some vesicular protein trafficking-associated genes, such as *TGOLN2* and *TMED2* (Fig. 2g, Supplementary Data 11). Ductal cells exclusively expressed additional trafficking genes including *RAB3IP, LAPTM4A, VPS37B*, and *TOR1A* as well as *SERPINA3* and *SERPING1*, which encode serine protease inhibitors (Fig. 2g, Supplementary Data 11).

At 24 hr, almost all autophagy-associated genes were upregulated, except for *FTL*, and were almost exclusively expressed in either β cells or monocytes (Extended Data Fig. 6d, Supplementary Data 11). We observed β cells upregulate several genes encoding lysosomal enzymes and compartments—*HEXB, PPT1, LAMP1, PSAP, ASAH1*, and *SCARB2*—and autophagy regulators—*DEPP1* and *WIPI1* (Extended Data Fig. 6d, Supplementary Data 11). We also noted positive expression of pro-apoptotic markers including *BNIP3* and *SH3GLB1* exclusively in β cells. α and β cells both upregulated *SQSTM1, VMP1, IDS*, and *NEU1*—all of which are involved in autophagy and lysosomal activity—as well as *FKBP8*, which is another pro-apoptotic marker (Extended Data Fig. 6d, Supplementary Data 11). On the other hand, most monocyte genes in the autophagy GO theme are associated with class II HLA expression and presentation. There was also increased expression of *FUCA1, CTSC, LGMN, NPC2*, and *CD68*, which are all genes encoding lysosomal components and proteases (Extended Data Fig. 6d, Supplementary Data 11). However, by 48 hr, the monocytes no longer differentially expressed any autophagy-related genes (Extended Data Fig. 6d).

Our data also showed downregulation of genes unique to β cells by 48 hr. Several of these in the vesicle GO theme—*HSPA1A, HSPA1B, CALR, PCMT1, GOLGA2*, and *KDELR1*—are involved in protein folding and transport (Fig. 2g, Supplementary Data 11). There was also decreased expression of multiple components of the vacuolar ATPase including *ATP6V0B, ATP6V0C*, and *ATP6V0E2* as well as proteinase inhibitors, such as *SERPINB6, TIMP2*, and *ITM2B* (Fig. 2g, Supplementary Data 11). In addition, autophagy- and lysosomal-associated genes, such as *TMEM59, WDR6, DNASE2*, and *GNPTG* were all downregulated by 48 hr in β cells (Extended Data Fig. 6d, Supplementary Data 11). Within the genes shared between the 24 hr and 48 hr autophagy gene lists in β cells, *ATP6AP1, ATP6V0C, EEF1A1*, and *PSAP* were upregulated 24 hr post infection but downregulated after 48 hr. β cells also had decreased expression of other vacuolar ATPase components, namely *ATP6V0B* and *ATP6V0E2* (Supplementary Data 11).

By 48 hr post infection, α cells demonstrated a robust response in vesicle-associated genes. Alongside ductal cells, α cells upregulated multiple proteosome subunits—*PSME2, PSMA6, PSMB8, PSMB10*, and *PSMA1* (Fig. 2g, Supplementary Data 11). Our data suggest α cells also increased expression of genes associated with protein and small molecule trafficking, such as *CD63, SLC36A4, DYNLT1*, and *RAB3B*, as well as cysteine proteinases including *CTSZ* and *CTSB*. We observed α cells overexpressed *VLDLR* and *LDLR*, which are involved in receptor-mediated endocytosis of low-density lipoproteins, while downregulating *SREBF1*—a transcription factor for genes involved in sterol biosynthesis (Supplementary Data 11). There was also downregulation of *NPC2*, a mediator of cholesterol transport through the lysosomal system. At 48 hr, α cells also downregulated multiple lysosomal/endosomal and vesicle-associated genes including *VAMP2, LAMP1, TOLLIP, SGSM1, STX3*, and *PSAP* (Fig. 2g, Supplementary Data 11).

Similar to the vesicle GO theme, α cells had a more robust autophagy-related response at 48 hr. We observed downregulation of a series of protein degradation-associated genes unique to α cells, such as *LAMP1, CTSD*, and *GGA2* (Extended Data Fig. 6d, Supplementary Data 11). The lysosomal membrane glycoprotein encoding gene, *LAMP2*, was also down in both α and mesenchymal cells. The mesenchyme underwent decreased expression of multiple lysosome-associated genes, such as *LITAF, KXD1*, and *LAPTM4A*. Components of the PI3K-mTORC signaling pathway, such as *PIK3R2, AKT1, PTEN*, and *LAMTOR4*, alongside translation-associated genes *EEF1A1* and *EEF1A2* were also downregulated 48 hr post infection in α, β, and mesenchymal cells (Extended Data Fig. 6d, Supplementary Data 11).

### CVB3 infection impacts the mitochondria in cell-specific manners

The mitochondria GO theme was the only theme associated with an organelle. Considering the importance of mitochondria to cell function, especially highly metabolic cells like pancreatic endocrine cells, we looked further into the genes within the theme. By 24 hr post infection, we observed β cells differentially regulated multiple genes associated with oxidative phosphorylation in mitochondria (Fig. 3a, Supplementary Data 11). CVB3-infected β cells differentially regulated multiple genes encoding mitochondrial complex I components. For instance, *NDUFS5, NDUFB7, NDUFA13*, and *NDUFA1* were all downregulated while *NDUFA6* and *NDUFA4* were upregulated. *SDHC*, a subunit of mitochondrial complex II, was upregulated. There were also changes in mitochondrial complex IV-associated genes where we noted *COX5B* and *COX16* were downregulated while *CYCS, COX7A1, COX11*, and *COX7B* were overexpressed. *ATP5MC2* and *ATP5PO*, which encode subunits of mitochondrial complex V, as well as *ETFDH* and *UQCRH*, which are associated with the electron-transport chain (ETC), were upregulated (Supplementary Data 11). Additionally, β cells underwent downregulation of genes involved in p53-mediated cell cycle arrest, including *CDKN1C, CDKN1A*, and *GADD45GIP1*, and upregulated a cyclin encoding gene, *CCNI*. There was also overexpression of antioxidant encoding genes—*GPX3, PRDX3*, and *GPX4*—and genes involved in molecule transport in/out of the mitochondria—*SLC25A3* and *SLC25A5*—in β cells (Supplementary Data 11).

**Fig. 3.**
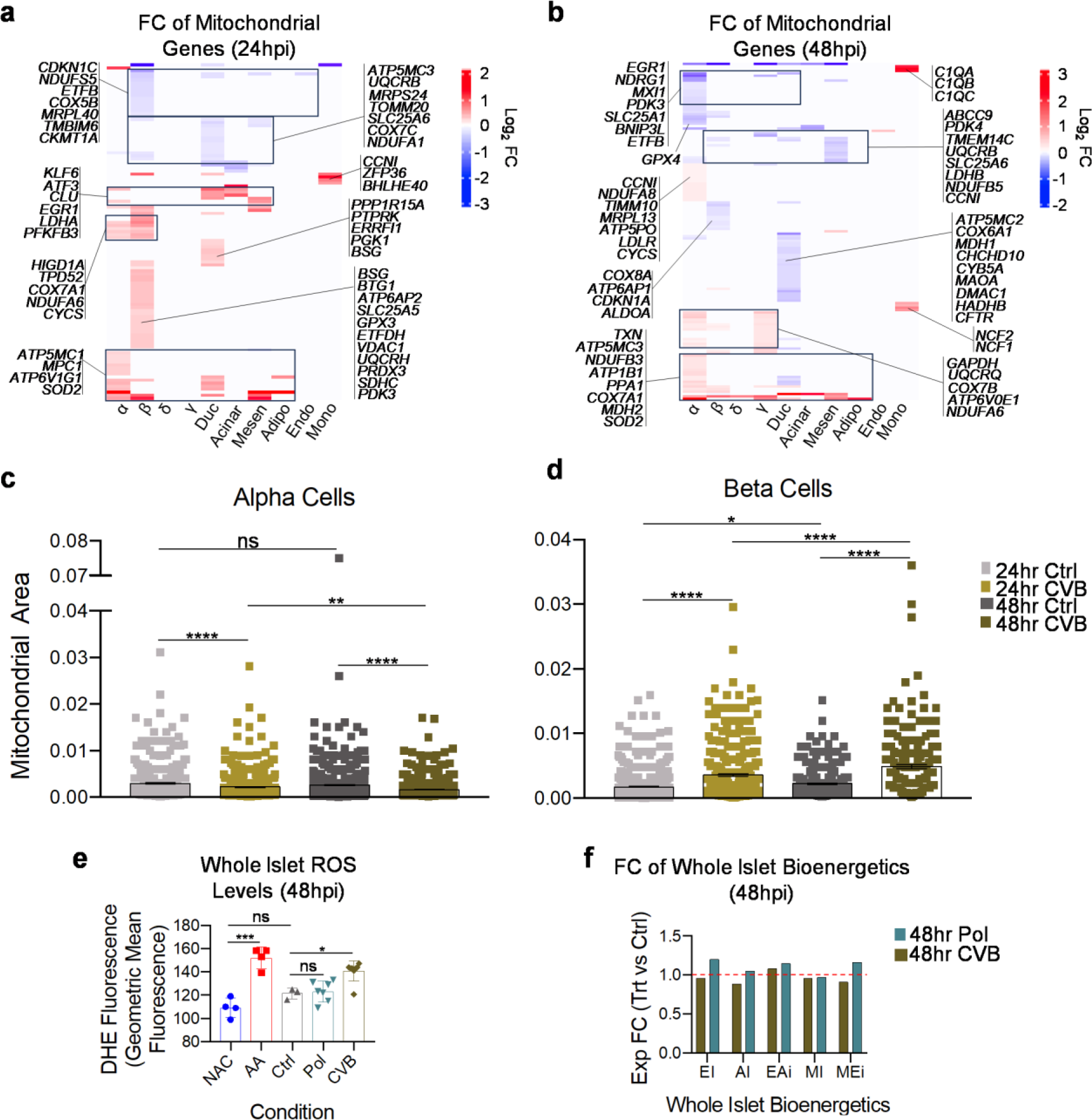
CVB3 infection induces mitochondrial dysfunction in primary human islets. **a, b**, Heatmaps depicting log_2_ fold change (FC) of mitochondria-associated genes across cell types when comparing each CVB3 timepoint to its respective control. **c**, Mitochondrial area of α cells based off measurements made from TEM images using ImageJ. (24 hr Ctrl, n = 494; 24 hr CVB3, n = 632; 48 hr Ctrl, n = 668; 48 hr CVB3, n = 815) ** = One-way ANOVA with Tukey’s multiple comparisons. Error bars represent the s.e.m. **d**, Mitochondrial area of β cells based off measurements made from TEM images using ImageJ (24 hr Ctrl, n = 669; 24 hr CVB3, n = 476; 48 hr Ctrl, n = 357; 48 hr CVB3, n = 210). ** = One-way ANOVA with Tukey’s multiple comparisons. Error bars represent the s.e.m. **e**, Dihydroethidium (DHE) fluorescence assay measuring reactive oxygen species production in whole primary islets 48 hr (NAC = N-acetyl Cysteine; n = 4. AA = Antimycin A; n = 4. Ctrl; n = 3. Pol; n = 7. CVB; n = 8). ** = One-way ANOVA with Tukey’s multiple comparisons. Error bars represent geometric s.d. **f**, Exponential FC of bioenergetics comparing poly(I:C) and CVB3 to Control at 48 hr. (n = 3. EI = ETC-dependent OCR proportion, AI = ATPase-dependent OCR proportion, EAi = ETC-dependent proportion of ATPase-independent OCR, MI = Maximal over initial OCR FC, MEi = Maximal over ETC-independent OCR FC). ** (Student’s t-test).

α cells also demonstrated differential regulation of multiple ETC components. We noted upregulated mitochondrial complex I genes—*NDUFA6* and *NDUFV2*—complex IV genes—*COX7A1* and *CYCS*—as well as *ATP5MC1*, a complex V gene, by 24 hr (Fig. 3a, Supplementary Data 11). We observed α cells increase expression of multiple glycolysis-associated genes, including *PDK4, LDHA, TPI1, ENO1, MPC1*, and *GAPDH. SOD2*, an antioxidant enzyme encoding gene, was also upregulated in α cells as well as mesenchyme and adipocytes at this timepoint (Fig. 3a, Supplementary Data 11).

Ductal cells downregulated *NDUFS5, NDUFB4*, and *NDUFA1* for mitochondrial complex I, *UQCRB* for complex III, *COX6C, COX7A2*, and *COX7C* for complex IV, and *ATP5MC3* and *ATP5MG* for complex V. *SLC25A6*, another gene involved in molecule transport in/out of the mitochondria, was also downregulated at 24 hr in ductal cells (Fig. 3a, Supplementary Data 11). This population upregulated *CCNI, EGR1*, and *DUSP1*, which are proliferation-associated genes.

The transcriptional response of acinar cells included downregulation of *AKR1C1* and *AKR1C3*, which both encode aldo-keto reductases, as well as *TXN*, a redox enzyme encoding gene. On the other hand, acinar cells upregulated *IFI6, KLF6, ATF3*, and *CLU* (Fig. 3a, Supplementary Data 11). Mesenchymal cells decreased expression of *NDUFS5, UQCRB*, and *ATP5MC3*, which are genes encoding subunits for mitochondrial complexes I, III, and V, respectively, while upregulating the glycolysis-associated genes, *LDHA, PFKFB3*, and *TPI1* (Fig. 3a, Supplementary Data 11).

At 48 hr, α cells had the strongest mitochondria-associated transcriptional response (Fig. 3b, Supplementary Data 11). These cells exhibited decreased gene expression for solute carrier proteins (*SLC25A5* and *SLC25A1*), components of the ETC (*ETFB* and *ETFDH*), apoptotic regulators (*NDRG1* and *BNIP3L*), and proliferation (*EGR1, CDKN1C, DUSP1*, and *CCNI*). α cells continued to exhibit increased expression of genes associated with mitochondrial complex I (*NDUFA8, NDUFA9, NDUFA1, NDUFB4, NDUFA12, NDUFB2, NDUFV2, NDUFA13, NDUFB1, NDUFAB1, NDUFC2*, and *NDUFB3*), complex III (*UQCRQ*), complex IV (*CYCS, COX5A, COX20, COX7A2, COX7B, COX14, COX17*, and *COX7A1*), and complex V (*ATP5PO, ATP5PD, ATP5MF, ATP5MC3*, and *ATP5MC1*) (Fig. 3b, Supplementary Data 11). α cells also differentially expressed apoptotic-associated genes, including *NDRG1, BNIP3L, IFI27L2, HIGD1A*, and *IER3* (Fig. 3b, Supplementary Data 11).

On the other hand, β cells did not exhibit unilateral expression of ETC-associated genes. β cells downregulated a complex III subunit (*UQCRB*), *COX8A* and *COX5B*, which are complex IV subunits, and a complex V subunit encoding gene, *ATP5F1D* (Fig. 3b, Supplementary Data 11). We also noted upregulation of the complex I subunit, *NDUFA4*, two complex IV-associated genes (*COX7B* and *COX7A1*), and a complex V subunit, *ATP5F1E*. Other mitochondria-associated genes in β cells included *MDH2*, which is important for oxidative phosphorylation, as well as downregulation of *TMEM14C* and *ALDOA*, which are involved in mitochondrial transport and glycolysis, respectively (Fig. 3b, Supplementary Data 11). We also observed decreased expression of the multiple components of vacuolar ATPase previously mentioned.

γ cells downregulated *FXYD2* and *ABCC9*, which encode transport channels, as well as a glycolysis regulator, *PDK4*. Yet, we observed upregulation of other glycolysis associated genes, *GAPDH* and *ENO1* (Fig. 3b, Supplementary Data 11). γ cells also increased the expression of mitochondrial complex I genes (*NDUFV2, NDUFA6, NDUFA13*, and *NDUFB1*), a subunit of complex III (*UQCRQ*), complex IV-associated genes (*COX7A2, COX7B*, and *COX14*), and complex V genes (*ATP5MF* and *ATP5F1E*).

Ductal cells mostly downregulated genes by 48 hr. Similarly to the other cell types, we observed decreased expression in genes associated with mitochondrial complex I (*NDUFA2, DMAC1, NDUFS1, NDUFA4*, and *NDUFA13*), complex III (*UQCRB*), complex IV (*COX8A, COX6A1, COX6C*, and *COX6B1*), and complex V (*ATP5MC2* and *ATP5F1C*). Ductal cells also decreased *COA1* expression, which is involved in complex I and IV assembly (Fig. 3b, Supplementary Data 11). Other notable mitochondrial function-associated genes with decreased expression in ductal cells included *TMEM14C, HMGB1, MDH1, CHCHD10, CYB5A, NNT, STMP1, MPC1, GATM*, and *HADHB* (Fig. 3b, Supplementary Data 11). Ductal cells increased expression of three genes only: *PPA1, IER3*, and *SOD2*. In acinar cells, we observed downregulation of *ATF3* and *FOS* and upregulation of *IFI6*.

Mesenchymal cells exhibited decreased expression of mitochondrial solute carriers (*SLC25A5, SLC25A6*, and *SLC25A3*), the *NDUFB5* complex I subunit, complex III subunits (*UQCRB* and *UQCRH*), as well as *TMEM14C* and *LDHB*, both of which are associated with mitochondrial function (Fig. 3b, Supplementary Data 11). On the other hand, mesenchymal cells upregulated apoptosis-associated genes (*TNFAIP3, IFI6*, and *IER3*) as well as ROS regulator, *SOD2*. Monocytes underwent increased expression of multiple polypeptides of the serum complement subcomponent C1q (*C1QA, C1QB*, and *C1QC*), NADPH subunits (*NCF1* and *NCF2*), and *TNFRSF1B*, which is an apoptosis regulator (Fig. 3b, Supplementary Data 11).

Considering that α and β cells had the most unique differentially expressed genes, 87 and 81, respectively, across both timepoints in the mitochondrial GO theme, we delved further into the effects of CVB3 infection on the mitochondria of these cell types. We utilized electron microscopy and took images of α and β cells infected with CVB3-eGFP to assess mitochondrial morphology (Extended Data 7a-h). We measured the mitochondrial size of these cell types and found the mitochondria in α cells decreased in area size in a time-dependent manner while mitochondria increased in area size throughout the same time in β cells when compared to respective controls (Fig. 3c, Supplementary Data 12). When comparing the mitochondria of α and β cells to each other, we observed smaller mitochondria in control β cells that is statistically significant at 24 hr but not at 48 hr, albeit with a similar trend in size difference (Extended Data Fig. 8a-b, Supplementary Data 12). However, β cell mitochondrial area size was significantly larger when compared to α cells at both timepoints upon CVB3-eGFP infection (Extended Data Fig. 8a-b).

We also investigated the impacts of CVB3 infection on mitochondrial function. To measure ROS production, we performed a dihydroethidium fluorescence assay (Supplementary Data 1). Control whole islets were either left untreated or cultured with either N-acetyl Cysteine, which functions as an antioxidant, or antimycin A (AA) to increase ROS levels. At 48 hr, infected islets had a significant increase in ROS production when compared to untreated islets (Fig. 3e).

Moreover, we treated whole islets with either poly(I:C) or CVB3-eGFP for 48 hr and then measured mitochondrial respiration after injection with oligomycin, FCCP, and AA/rotenone using the Seahorse XF system (Supplementary Data 13). We employed the methodology found in Yépez et. al to perform oxygen consumption rates (OCR) statistics on the results.^30^ Briefly, we applied linear regression across the dataset to define intervals corresponding to each injection treatment, allowing us to model OCR changes. To compare the mitochondrial function in treated versus control islets, we computed bioenergetic values by taking the natural log difference of two corresponding OCR interval estimates. This log difference was then exponentiated to express the fold change between treatment conditions. To determine statistical significance, we applied linear regression analysis to these fold changes and conducted a Student’s t-test, generating p-values for each bioenergetic comparison to quantify the probability of the observed effects under the null hypothesis.

At 24 hr, poly(I:C)-treated whole islets exhibited lower values for ETC-dependent OCR proportion (EI), the ATPase-dependent OCR proportion (AI), the ETC-dependent proportion of ATPase independent OCR (EAi), and the maximal over ETC-independent OCR proportion (MEi) while the maximal over initial OCR proportion (MI) was higher relative to control islets (Extended Data Fig. 8c-d). Whole islets infected with CVB3 for 24 hr showed increases in MI, MEi, and EAi and decreased values for AI and EI (Extended Data Fig. 8c-d). For poly(I:C)-treated islets compared to control at 48 hr, we observed positive exponential fold changes for EI, AI, EAi, and MEi while MI decreased (Fig. 3f, Extended Data Fig. 8e-f). For whole islets infected with CVB3 for 48 hr, the EI, AI, MI, and MEi all decreased relative to control while the EAi increased (Fig. 3f, Extended Data Fig. 8e-f).

### Trajectory analysis reveals progressive transcriptional changes in response to CVB3 infection

After a viral infection, cells can display a spectrum of dynamic molecular states, each yielding distinct functional consequences. Thus, to delineate transcriptional differences between treatments, we independently reclustered β, α, and ductal cells, which respectively generated four, four, and three subpopulations (Fig. 4a-b, Extended Data Fig. 9a-d). To help capture intermediate cell states and elucidate the relative progression of CVB3 infection, we employed recent advances in objectively choosing the earliest principal node, which was characterized by an area with high number of control cells, and conducted pseudotime trajectory analysis using Monocle3.^31–34^ Our trajectory analysis revealed β3 and β4 were the most transcriptionally different to β1 and progressively increased expression of immune response genes (*PSMA7, CEBPB*, and *UBXN1)*, DNA damage-associated genes (*PHF1, CDKN2AIP*, and *JUND*), autophagy genes (*MAP1LC3B* and *MT2A*), and apoptosis-related genes (*ATF4, KLF11, DDIT3*, and *TRIB3*) (Fig. 4c-d, Supplementary Data 14).

**Fig. 4.**
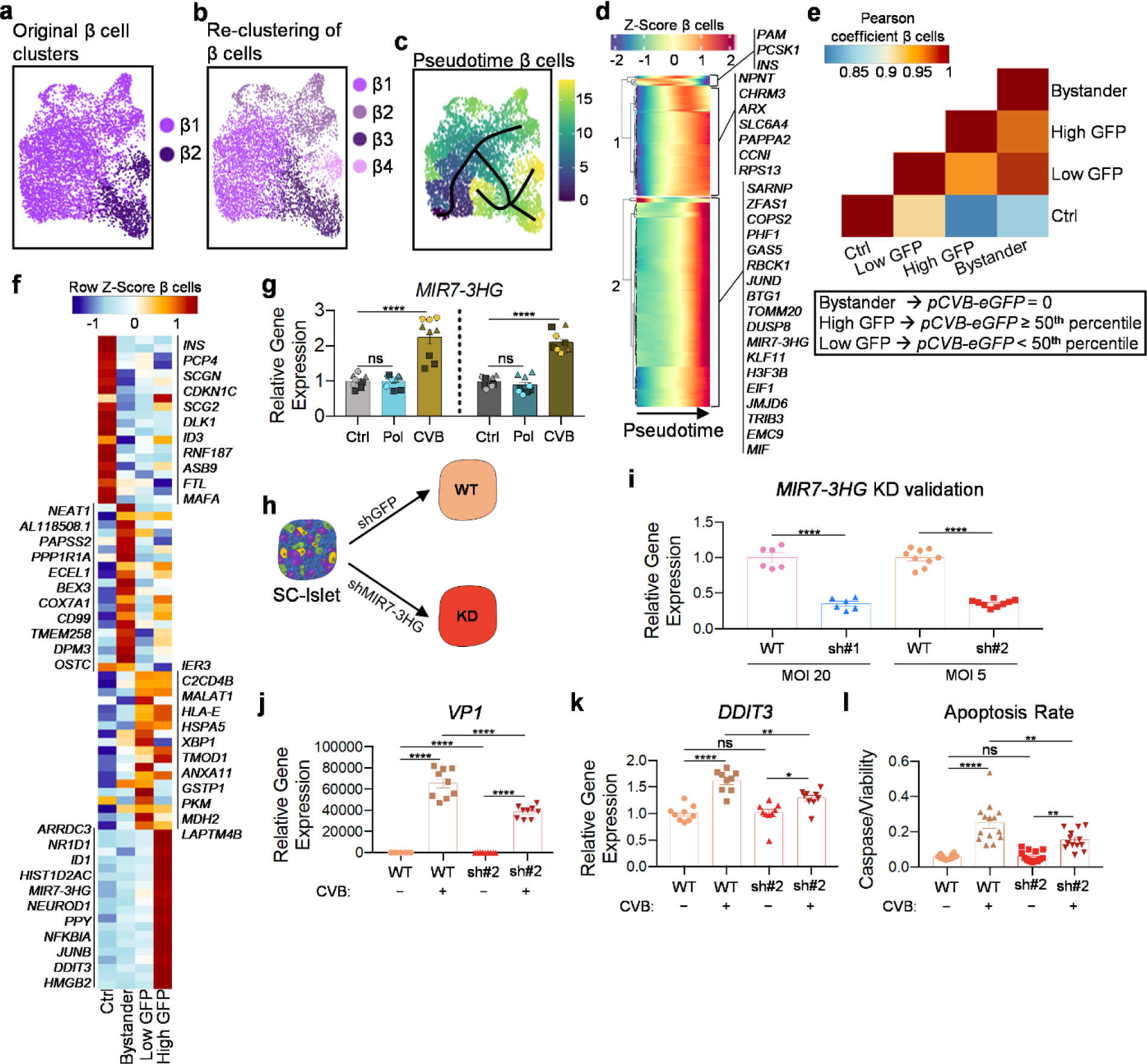
β cells exhibit differential transcriptional responses dependent on viral load that help identify *MIR7-3HG* as an apoptosis regulator in stem cell-derived islets infected with CVB3. **a**, UMAP showing original distribution of β cell populations (subset of β cell populations from all conditions). **b**, UMAP showing subpopulations upon re-clustering of β cell populations only (subset of β cell populations from all conditions). **c**, UMAP showing the pseudotime trajectory of β cell subpopulations (subset of β cell populations from Control and CVB3 conditions). **d**, Pseudotime trajectory heatmap showing the dynamic changes in β cell gene expression upon CVB3 infection. **e**, Pearson correlation plot of β cell subpopulations grouped by viral load (subset of β cell populations from Control and CVB3 conditions). **f**, Heatmap of the top upregulated genes in control, bystander, low GFP, and high GFP β cells. **g**, rt-qPCR of *MIR7-3HG* using RNA extracted from scRNA-seq patients (n = 3. Patient 01 = Triangle, Patient 02 = Square, Patient 03 = Circle). **h**, Schematic of experimental set up to knock down *MIR7-3HG* in SC-islets. **i**, rt-qPCR showing knockdown of *MIR7-3HG* with two different short-hairpin RNA vectors (sh#1; n = 6. sh#2; n = 9). **j**, rt-qPCR of viral antigen gene expression upon knockdown of *MIR7-3HG* and CVB3 infection of SC-islets (n = 9). **k**, rt-qPCR of apoptosis marker gene expression upon knockdown of *MIR7-3HG* and CVB3 infection of SC-islets (n = 9). **l**, Apoptosis assay measuring caspase 3/7 activity normalized to cell viability (n = 14). ** One-Way ANOVA. Error bars represent the s.e.m.

Upon further investigation, we also observed higher expression levels of multiple genes associated with transcription along the β cell-pseudotime trajectory. Some of these genes included the transcriptional regulators *ARX*, and *CCNL1*, chromatin remodelers, including *AKAP8L* and *HMGB2*, as well as proteins directly involved in the transcriptional process: *BTF3, TCEA1, TAF1D, POLR1D, POLR2H*, and *GTF3A* (Fig. 4d, Supplementary Data 14). β3 and β4 cells also highly expressed mRNA splicing factors—*SNRPG, SRSF7, SRSF3, CCNL1*, and *CLK1—*and translation-associated genes, such as multiple Eukaryotic Initiation Factors (EIFs), *SARS, CNBP*, and *MKNK2* (Fig. 4d, Supplementary Data 14). These cells expressed multiple regulators of circadian rhythm (*NR1D2, RACK1, PER1*, and *BHLHE41*), and proliferation-associated genes (*TPT1, BTG1, GGNBP2*, and *SYF2*). A multitude of long non-coding RNAs, including several small nucleolar RNA host genes (SNHGs), *GPC-AS1, LINC01578*, and *MIR7-3HG* were also overexpressed (Fig. 4d, Supplementary Data 14).

In α cells, α4 was the most transcriptionally different from α2 (Extended Data Fig. 9e). α cells also progressively downregulated cell identity and function genes (*MAFB, CHGB, PCSK2*, and *PDK4*) and upregulated mitochondrial and antiviral response genes as previously reported (Extended Data Fig. 9i, Supplementary Data 14). Nevertheless, ductal cells 1 (Duc1) were the most transcriptionally different from Duc2 in our trajectory analysis (Extended Data Fig. 9g). Ductal cells were predicted to progressively increase genes associated with cell cycle regulation and proliferation (*RACK1, S100A10, RASL12, CCND1*, and *CDKN2B*), mitochondria, ribosomal proteins and translation (*RPL7A, EEF1A1, SNHG6, NACA*, and *TRMT112*), the cytoskeleton and cell adhesion (*TMSB10, CFL1, TPM1, MYL12A, TUBA1A, TMSB4X, ACTB, ITGB1*, and *LAMB3*), and the ECM (*MMP7, LUM, COL4A2, BGN, DCN*, and *OLFML1*) (Extended Data Fig. 9k, Supplementary Data 14).

Next, we sought to determine the impacts of viral presence on transcriptional signatures by measuring levels of viral transcripts. To achieve this, we leveraged the *eGFP* transcript fused to the CVB3 viral genome. Briefly, we subsetted CVB3-infected cells and calculated the gene expression value of *pCVB3-eGFP* corresponding to the 50^th^ percentile and divided the cells into three groups: Bystander, Low GFP, and High GFP. The Bystander group was composed of cells with no *pCVB3-eGFP* expression, Low GFP cells were those with *pCVB3-eGFP* expression values less than the 50^th^ percentile, and High GFP cells expressed this transcript at a value greater than or equal to the 50^th^ percentile. We also included the corresponding control cells in our downstream analyses.

In β cells, Pearson correlation analysis revealed Low GFP and Bystander cells were the most transcriptionally similar and both were the most dissimilar to the control group (Fig. 4e). High GFP cells were mostly correlated with Bystander cells as well. Control α cells were most correlated with bystander cells followed by low GFP α cells (Extended Data Fig. 9f). α cells with high GFP were most like low GFP α cells. On the other hand, bystander ductal cells were least correlated with control and high GFP groups while low and high GFP cells were correlated (Extended Data Fig. 9h).

We then identified differentially expressed genes between these groups and graphed the genes with highest expression (Supplementary Data 14). Overall, control β cells had higher expression of genes associated with calcium binding (*PCP4* and *SCGN*), E3 ubiquitin-protein ligases (*RNF187* and *ASB9*), and β cell identity (*INS, DLK1*, and *MAFA*) (Fig. 4f). Whereas *HOPX, HSPB6, TMEM258, DPM3*, and *OSTC*, which are all involved in protein folding, were upregulated in bystander β cells alongside *NEAT1* and *AL118508*.*1*, which are both lncRNAs (Fig. 4f, Supplementary Data 14). Bystander β cells also overexpressed cell death-associated genes *BEX3* and *CD99* as well as *PAPSS2*, which is a 3’-phosphoadenosine 5’-phosphosulfate synthetase (Fig. 4f). Low GFP β cells overexpressed the cell cycle regulator *MALAT1, CHGA, HSPA5*, and *ANXA11*, which is an annexin protein (Fig. 4f, Supplementary Data 14). Both low and high GFP β cells expressed stress response genes, including *IER3, HLA-E, DNAJC12*, and *XBP1* as well as cytoskeleton-associated genes (*ACTG1* and *TMOD1*) and calcium signaling genes (*C2CD4A* and *C2CD4B*). High GFP cells overexpressed unique genes, such as *ARRDC3, NR1D1, NEUROD1, JUN, DDIT3*, and *MT-ND4L* (Fig. 4f, Supplementary Data 14). These cells also expressed transcriptional regulators and mRNA-decay associated genes (*ZFP36L2, HNRNPDL*, and *JUNB*) (Fig. 4f, Supplementary Data 14).

Control α cells expressed many genes involved in mitochondrial function (*HMGCS2 EPHX1, SLC7A2, NDRG1*, and *SLC25A5*), trafficking (*ARRDC4* and *S100A10*), lipid metabolism (*PDZK1, PLIN3*), and proliferation (*MOB1B, EGR1, PLCE1*, and *ERBB3*) (Extended Data Fig. 9j, Supplementary Data 14). We mostly observed overexpression of oxidative stress, including *MT2A, COX6C, NDUFB1*, and *MT-ND4L*, and lipid metabolism (*DBI*) in bystander α cells. Low GFP α cells upregulated many immune and stress response genes (*CEBPD, IFITM2, SOCS1, XBP1, NFKBIA*, and *BIRC3*) alongside calcium signaling genes (*C2CD4A* and *C2CD4B*), cytoskeletal genes (*TUBA1A* and *VIM*), and the autophagy associated *VMP1* (Extended Data Fig. 9j, Supplementary Data 14). High GFP α cells overexpressed multiple non-coding RNAs (*MALAT1, MIR7-3HG, NEAT1*, and *AC099509*.*1*) and stress response genes (*ELF3, INHBA, MUC13*, and *HPSB1*).

Control ductal cells expressed multiple signaling genes (*FOSB, IMPA2*, and *APCDD1*), transporters (*SLC3A1* and *SLC4A4*), and mitochondrial genes (*ATP5MC3, NDUFA4, COX6C*, and *ATP5MC1*) (Extended Data Fig. 9l, Supplementary Data 14). The top positively regulated genes for all non-control ductal cells were mostly linked to immune response and inflammation. Genes expressed in bystander ductal cells were not highly expressed in virus infected cells (*IAPP, SAA2, HLA-DRB5, SAA1, LCN2, HLA-DQB1*, and *OCIAD2*). Many of the immune-associated genes upregulated in High GFP ductal cells were also expressed in low viral titer cells, including *IFI44L, OAS3, ADAR, ISG15, MX1, STAT1*, and *TAP1* (Extended Data Fig. 9l, Supplementary Data 14).

### MIR7-3HG regulates viral amounts and response in stem cell-derived pancreatic islets

To alleviate the effects of CVB3 infection, we explored novel, antiviral response-specific genes. Our analysis identified *MIR7-3HG* as an endocrine-specific lncRNA that was only present in virally infected cells and within β cells, it was expressed highest in cells with high viral titer (Fig. 4f-g, Supplementary Data 1, Supplementary Data 14). *MIR7-3HG* has not been studied in the context of human primary islets. In cancer cell lines, it serves as a potent inhibitor of autophagy and part of a positive feedback loop involving its transcriptional regulator, MYC.^35^ To decipher the role *MIR7-3HG* plays in endocrine antiviral responses, we utilized short hairpin RNA (shRNAs) lentiviral vectors to knock down this gene in stem cell-derived pancreatic islets (SC-islets) (Fig. 4h).

Our lab utilizes a six-stage protocol that mimics pancreatic development and differentiates human pluripotent stem cells (hPSCs) into SC-islets by regulating various signaling pathways using small molecules and proteins (Extended Data Fig. 10a, Supplementary Data 1).^36,37^ These cells are more than 80% endocrine and capable of secreting insulin following glucose stimulation. In addition, our SC-islets are capable of restoring normoglycemia two weeks after transplantation into diabetic NOD mice.^36–38^ SC-islets can be generated in abundance, have the same genetic background across batches, and provide a human specific model. These characteristics allowed us to replicate and perform robust experiments.

We achieved a 65% knock down with shRNA #1 and 69% with shRNA #2 that was maintained upon CVB3 infection of HUES8 SC-islets (Fig. 4i). Upon *MIR7-3HG* knockdown, we observed via rt-qPCR a decrease in the relative expression of the viral gene *VP1* following CVB3 infection (Fig. 4j, Extended Data Fig. 10b, Supplementary Data 1). Considering this, we next sought to investigate the effects of *MIR7-3HG* knockdown on antiviral response in SC-islets and found there was an overall trend for downregulation of *IRF1, DDX58, IFIH1*, and *ISG15* (Extended Data Fig. 10c-f). We did observe a significant decrease in *IRF1* in SC-islets treated with shRNA #1 (Extended Data Fig. 10c). We also looked at islet identity genes and saw relatively no effect (Extended Data Fig. 10g-k).

Despite that, we found there was a significant reduction in the relative expression of the pro-apoptotic gene *DDIT3* upon CVB3 infection (Fig. 4k, Supplementary Data 1). We followed up on this observation using shRNA #2 since it obtained a desirable knock down with a lower multiplicity of infection. To validate our gene expression assays, we multiplexed the CellTiter-Fluor Cell Viability Assay with the Caspase-Glo 3/7 Assay System on plated down SC-islets (Supplementary Data 1). Following CVB3 infection, *MIR7-3HG* knock down decreased the apoptosis rate in CVB3-infected SC-islets compared to infected controls. However, the apoptosis rate was still higher when compared to non-infected controls (Fig. 4l).

## Discussion

Environmental factors, such as viruses, are closely associated with the induction and progression of T1D. Despite that, the relationship between CVB infection and T1D remains unclear, largely due to limitations in current technologies and animal models. An increased understanding of pancreatic islet response to CVB infection could improve therapies to prevent or delay T1D following viral infection. However, delineating cell-type specific responses in primary human tissues has proven difficult due to the heterogenous cell composition found within islets. In the present study, we used scRNA-seq to provide a more robust representation of how islets respond to CVB3 infection over time while highlighting pathways and genes associated with viral stress in different cell types. Comparison with poly(I:C)—a synthetic double-stranded RNA—identified the limitations of this control treatment since it does not mimic the transcriptional complexity induced by viruses. Although our scRNA-seq dataset was limited to four cadaveric donors, it still provided important information for downstream functional assays that identified unique cell-specific effects and genetic candidates. The genes and cellular processes highlighted by our data can be targeted to alleviate the impacts of CVB infection on pancreatic islets.

Our analysis of whole islets revealed major global transcriptional differences between CVB3 infection and poly(I:C) treatment across two timepoints. Although there were similarities between treatments, CVB3 induced a far more complex transcriptional response supported by a much higher number of differentially expressed genes associated with a multitude of cell processes. At both timepoints, the CVB3-treated islets experienced changes in genes linked to calcium signaling, the ECM, ER stress, ubiquitin ligases, and transcription whereas poly(I:C) induced an immune response characterized mostly by increased expression of type 1 IFN-related genes and differential expression of various ribosomal proteins. Previous reports have shown that CVB3-encoded proteins are involved in more than just viral genome replication in HeLa cells and can affect cellular processes, such as protein trafficking and autophagy, to evade the immune system and enhance its replication.^21,26,39^ This highlights the importance of studying the effects of viral infection beyond immune response in studying its connection to T1D pathogenesis.

In this study, we leveraged scRNA-seq to delineate the cell-type specific effects of CVB3 infection on pancreatic islets. Until now, CVB has been shown to infect human ductal and β cells^23–25,40–42^ with little evidence on how it targets or impacts other pancreatic cell types in humans.^43^ In this study, we demonstrated CVB3 infects all islet cell types at similar levels despite very low mean gene co-expression of both entry receptors in all populations except δ and ductal cells. Although detection of CD55 and CAR co-expression is a standard method to measure CVB tissue permissiveness,^15^ there have been reports of CVB detection in cells not expressing CAR, including human β cells.^15,23–25,28,42,44–47^ Although the entry mechanisms used for CVB remain unknown, our dataset proves useful in investigating this in the future.

A comparison of each cell type to its respective control showed β, α, and ductal cells were the most impacted and each had unique transcriptional responses following CVB3 infection. α and ductal cells had the strongest type 1 IFN-associated response and maintained increased expression of immune response genes at 48 hr. In α cells, our scRNA-seq data suggested CVB3 also upregulates proteasome components and affects expression of genes associated with lipid biology. However, by 24 hr, β cells predominantly upregulated genes related to autophagy and lysosomal protein degradation, while by 48 hr, there was a reduction in the expression of genes associated with autophagy, as well as those involved in protein folding and transport.

Autophagy can be both a positive and negative regulator of IFN signaling and can also promote pathogen clearance.^48^ Others have shown CVB and other enteroviruses induce cellular autophagy to disrupt selective degradation of viral proteins, disrupt NFκB signaling to evade the immune system, and promote viral replication and release.^39,49–53^ CVB non-structural proteins have been reported to reduce surface expression of B2M and class I HLAs as well as protein secretion by restricting anterograde trafficking in the Golgi Apparatus (Golgi), disrupting the Golgi complex, and increasing the rate of endocytosis.^21,26^ Viruses have also been known to reduce endogenous translation to preferentially drive viral protein translation.^54^ This may be plausible in β cells considering the increase in EIF genes as well as mRNA-decay genes we observe in infected cells. Thus, unlike other pancreatic cell types, β cells may serve as a main replication hub for CVB during pathogenesis.

Viral infections have been known to impact various aspects of the mitochondria.^55,56^ At the transcriptional level, we observed unique cell-type signatures with genes associated with mitochondria. At both timepoints, β cells differentially expressed ETC-associated genes and increased antioxidant gene expression while α and γ cells upregulated ETC-associated genes. α cells also increased glycolysis gene expression. Ductal cells underwent down regulation of ETC- and mitochondrial function-associated genes.

In the present study, we also reported mitochondria size was decreased in α cells and increased in β cells. Viruses can cause either mitochondrial fusion or fission but have not been reported to cause both.^55^ However, it remains unclear if these mechanisms are involved in pancreatic endocrine cells. We also validated our scRNA-seq data and showed CVB3 infection induces mitochondrial dysfunction and leads to an increase in ROS levels at the whole islet level. The observed changes in ETC-associated genes and increase in ROS levels could mark a metabolic reprogramming to aerobic glycolysis, which all together would provide the necessary energy for viral replication while avoiding further increases in ROS by diverting fuel away from oxidative phosphorylation.^54,57^ To keep the cell alive to support their infection cycle, multiple viruses increase antioxidant enzyme expression to regulate ROS levels caused by calcium signaling disruption, ER stress and more.^54^ CVB has also been shown to induce and alter mitophagy in murine cells to facilitate the release of virus-containing mitophagosomes.^58^ Therefore, our autophagy transcriptional data as well as our mitochondrial transcriptional and functional data demonstrate CVB3 could impact mitochondria to promote viral replication and/or release in multiple islet cell types.

Our data highlight multiple cellular processes that are impacted in pancreatic islets and serve as a resource for future investigations regarding the connection between T1D and CVB infection. Considering ductal cells can be persistently infected by CVB^40^ as well as our observation that they exhibited the strongest cytokine response among the various cell types, cross-tissue communication may serve as a potential trigger for T1D following persistent viral infection by contributing to the cytokine storm preceding disease onset. CVB3 also induced expression of multiple genes associated with T1D (*IER3, PTPRN2, ZFP36L1, HLA-DRB1, RASGRP1, POMC, IRF1, HLA-B*, and *TNFRSF11B*).^59–71^ Thus, persistent infection in genetically susceptible individuals could increase the risk of T1D initiation. CVB may also be linked to T1D through its effect on autophagy function. Cytokines have been shown to block autophagy flux and increase the amount of autophagosomes in β cells.^72^ A previous study reported impaired autolysosome formation in T1D mouse models and human T1D pancreatic tissue that led to reduced degradation by lysosomal enzymes.^73^ Thus, CVB may contribute to T1D through its disruption of autophagy flux to enhance its replication.^49,52^ β cells are also among the most metabolically active tissues in the body and CVB infection has been shown to negatively impact insulin secretion in human primary islets.^23,24^ Therefore, the detrimental effects of CVB3 infection on mitochondrial function, ETC gene expression, and the ER we observed may lead to T1D. Most cells rely on antioxidant enzymes to return ROS levels to normal.^13^ However, human and rodent β cells have unusually low levels of antioxidant enzyme activity, which renders them sensitive to oxidative stress.^74,75^ Elevated ROS levels and ER stress have been shown to contribute to impaired insulin secretion as well as reduction of insulin gene expression and content in β cells.^76–81^ Thus, the increased ROS levels, ER stress, and impaired function caused by CVB infection could contribute to T1D by inducing β cell death.

This study has some limitations and topics to be addressed in future studies. Although our comprehensive analysis serves as a valuable tool to improve our understanding of viral effects on pancreatic islets, our dataset could be expanded by the addition of more donors, especially T1D patients, to capture a better representation of the population. Expanding the cell number in our dataset could also capture valuable transcriptional information in poorly represented cell types, such as adipocytes. Incorporating single-cell/nuclei assay for transposase-accessible chromatin (ATAC) sequencing, which has been recently used to study and understand SC-islet differentiation and *CFTR* in type 1 diabetes, could provide further insights.^82–84^ Furthermore, single-cell sequencing analysis has been previously used to study primary islets, SC-islets, and primary islets under cytokine and small molecule stressors, and cross-referencing our new datasets to prior work could yield new insights.^82,83,85–91^

Here, we only focused on CVB3 infection at 24 hr and 48 hr. Future investigations could include longer timepoints and establish a model of persistent infection in islets as previously done in ductal cell lines.^40^ Further work must also be done to validate the observed transcriptional signatures, including changes in autophagy and CVB3 infection of all cell types. One possible explanation for infection of the whole islet could be viral transfer via vesicles or other enveloping cell structures.^28,53,58,92^

In the present study, we proved how our dataset could also be used to find novel candidate genes to alleviate the impacts of viral infection. We were the first to report that knockdown of the lncRNA *MIR7-3HG* in human SC-islets can reduce gene expression of the viral protein *VP1* and lowers apoptotic rates following CVB3 infection. Considering its role in negatively regulating autophagy in cancer cell lines, it is possible *MIR7-3HG* knockdown is activating the autophagy machinery and promoting viral protein degradation.^35^ This study also leveraged SC-islets as a model system for study, which has been used previously in many disease modeling systems,^93–99^ but is the first to be used to study CVB for diabetes. Overall, our study unveils novel cell-type specific responses to CVB infection in human primary islets that can be used in the future to improve antiviral and diabetes treatment.

## Author contributions

D.A.V-P. and J.R.M. designed all experiments and wrote the manuscript. D.A.V-P. performed all sequencing and computational experiments. D.A.V-P. and D.H. performed all *in vitro* experiments. J.P.T. and H.M.T. and provided key guidance, insights, and reagents. All authors revised, reviewed, and approved the manuscript.

## Acknowledgements

This work was primarily funded by the NIH (R01DK138469) to J.R.M. and H.M.T. Further support was provided by the NIH (R01DK114233, R01DK127497, R01DK126456), JDRF (3-SRA-2023-1295-S-B), a Human Islet Research Network (HIRN) Catalyst Award, the Edward J. Mallinckrodt Foundation, and startup funds from the Washington University School of Medicine Department of Medicine to J.R.M. Further support was provided by the Washington University Diabetes Research Center (P30DK020579). We thank the Genome Technology Access Center at the McDonnell Genome Institute at Washington University School of Medicine for help with genomic analysis. The Center is partially supported by NCI Cancer Center Support Grant P30CA91842 to the Siteman Cancer Center from the National Center for Research Resources (NCRR), a component of the National Institutes of Health (NIH), and NIH Roadmap for Medical Research. D.A.V-P. was supported by the NSF Graduate Research Fellowship Program (DGE-2139839 and DGE-1745038). This publication is solely the responsibility of the authors and does not necessarily represent the official view of NIH nor any other funder. We would also like to thank Erika Brown (Washington University) and Punn Augsornworawat (Mahidol University) for helpful feedback on the manuscript.

## Competing Interests

J.R.M. is an inventor on licensed patents and patent applications related to SC-islets. J.R.M. was employed at and has stock in Sana Biotechnology. The remaining authors declare no competing interests.

## Methods

### Approvals

Our research adheres to all pertinent ethical regulations. Non-diabetic islets were sourced from Prodo Laboratories Inc., which ensures informed consent for all non-identifiable information. No compensation is provided for these islets, which have been rejected for transplantation, quality-controlled, and deemed suitable for research purposes only. The study design did not consider sex as it was outside the scope of this research. Approval for this research was granted by the Washington University Institutional Biological & Chemical (IBC) Safety Committee (Approval number 12186). All work involving HUES8 was approved by the Washington University Embryonic Stem Cell Research Oversight Committee (Approval number 15-002).

### Culture of primary human islets

Human islets were obtained from Prodo Laboratories Inc. Upon arrival, they were spun down at 100 x *g* for one min and then resuspended and cultured in CMRL 1066 media supplemented (Corning, 99-603-CV) with 10% FBS. Islets were cultured in 6-well plates with 2,000 islet equivalents (IEQ) and 5 mL of media per well and in suspension on an Orbi-Shaker (Benchmark) set at 100 RPM. They were cultured in a humified incubator at 5% CO_2_ and 37 °C for at least 48 hr before treatment.

### Whole Islet Infection/Treatment

On infection day, the number of islet clusters per well was assessed via counting under a microscope. The average from at least two independent counts was calculated. Assuming there were 1,000 cells per cluster, the total number of cells was estimated by multiplying the number of islet clusters by 1,000. Islets were then treated with either endotoxin-free water, 500 ng/mL polyinosinic-polycytidylic acid [poly(I:C)] (InvivoGen, tlrl-piclv), or CVB3-eGFP (titer = 1.63 × 10^8^ pfu/mL) at MOI of 20. Cells were collected for downstream assays 24 hr or 48 hr post-treatment.

### Real-time quantitative PCR

Cells were collected, washed with 1X PBS (Corning, 21-030-CV), and used for RNA extraction using the RNeasy Mini Kit (Qiagen, 74106) with DNase treatment (Qiagen, 79254). Reverse transcription PCR reactions were performed with 100-200 ng total RNA using the High-Capacity cDNA Reverse Transcriptase Kit (Applied Biosystems, 4368813) and a T100 Thermocycler (BioRad). Real-time quantitative PCR reactions were then performed with PowerUp SYBR Green Master Mix (Life Technologies, A25742). A QuantStudio 6 Pro Thermocycler (Applied Biosystems) was used for all PCR reactions. Analyses were done using the ΔΔCt methodology. The average Ct of the *TBP* and *GUSB* housekeeping genes was used for normalization.

### Cell preparation for scRNA sequencing

On submission day, whole primary islets for patients 01-03 were washed with 1X PBS (Corning, 21-030-CV) and single cell dispersed with TrypLE Express (Thermo Fisher, 12604-039) at 37 °C for ∼16 min with a gentle swirl every 5 min. The dispersed cells were then washed three times with cold PBS and centrifuged at 300 x *g* for 3 min after each wash. TotalSeq™ antibody cocktails (BioLegend, 3946-01, -03, -05, - 07, -09, -11) were simultaneously prepared by adding 1 μg of each TotalSeq™ antibody to 300 μL of Cell Staining Buffer and centrifuged at 14,000 x *g* at 4 °C for 10 min. After careful aspiration, the cell pellet from each treatment condition and time point was resuspended in a TotalSeq™ antibody cocktail and incubated for 30 min at 4 °C. The cells were then washed 3 times with 1 mL of cold Cell Staining Buffer and centrifuged at 350 x *g* for 5 min at 4 °C after each wash. The cells were also carefully resuspended with a p200 micropipette with each wash. The cells were then resuspended in cold DMEM at 1,000 cells/μL. All different samples were then pooled at equal proportions. The cell concentration was verified using a Countess II (Invitrogen). All submissions had a cell concentration near 1,000 cells/μL and cell viability above 70% except one with 51%.

Whole primary islets from patient 04 underwent the same single cell dispersal and wash procedure. The cell pellet from each condition and timepoint were then resuspended in cold DMEM at 1,000 cells/μL and underwent the same verifications prior to submission. All submissions had a cell concentration near 1,000 cells/μL and cell viability above 70%. TotalSeq™ antibody ID and sequence:

1. Hashtag_TotalSeq_A0251— GTCAACTCTTTAGCG
2. Hashtag_TotalSeq_A0252— TGATGGCCTATTGGG
3. Hashtag_TotalSeq_A0253— TTCCGCCTCTCTTTG
4. Hashtag_TotalSeq_A0254— AGTAAGTTCAGCGTA
5. Hashtag_TotalSeq_A0255— AAGTATCGTTTCGCA
6. Hashtag_TotalSeq_A0256— GGTTGCCAGATGTCA

### scRNA-sequencing

The samples were submitted to Washington University McDonnell Genome Institute for library preparation and sequencing. The libraries were constructed using the Chromium Single Cell 3’ v3.1 Library and Gel Bead Kit (10X Genomics). The libraries were then sequenced using a NovaSeq6000 System (Illumina).

### scRNA sequencing analysis

#### Raw data processing

Seurat v4.3.0.1 was used on R Studio 1.3.1093 (R version 4.0.3) for single cell analyses and comparison between all groups.^100^ The samples were aligned to the human GRCh38 reference genome and then underwent quality control filtering. Although filtering parameters differed between conditions and patients, only cells with overall mitochondrial percentages less than 10-25%, a minimum gene number range of 500-3500, a maximum gene number range of 6000-9500, and a maximum number of molecules range of 30,000-150,000 were used for analysis. These ranges were chosen to remove any dying cells, debris, and doublets based on the corresponding distributions.

#### Dataset normalization, integration

Gene expression data were normalized to correct for batch differences using *SCTransform* in conjunction with the glmGamPoi package. Mitochondrial percentage variables were regressed. Repeatedly variable features and anchors were identified using the *SelectIntegrationFeatures* and *FindIntegrationAnchors* functions, respectively, before integration with *IntegrateData*.

#### Clustering

Dimensionality reduction was performed by PCA and UMAP embedding and cells were then plotted based on similar gene expression patterns using the *FindNeighbors* and *FindClusters* functions with a 0.4 resolution. *PrepSCTFindMarkers* followed by *FindAllMarkers* using the “wilcox” test were run to identify differentially expressed genes in each cluster compared to all other clusters, which were then used to designate the different cell types within primary human pancreatic islets. The *FeaturePlot* function was used to visualize the expression of specific genes across different cell types while *heatmap*.*2* within the gplots package created a heatmap depicting average expression values of marker genes for each cell type, which were obtained via the *AverageExpression* function. *DotPlot* was used to graph *MIR7-3HG* expression across cell types.

#### Analyses of Poly(I:C), CVB3, and Control whole islet conditions

The *subset* function was used to isolate the 24 hr and 48 hr timepoints. For each timepoint, *AverageExpression* was run on cell marker genes as well as type 1 interferon-associated genes and the results were plotted using the *heatmap*.*2* function. All possible pairwise comparisons between conditions within each timepoint were conducted using the *FindMarkers* function with no logfc.threshold. Differentially expressed genes with an adjusted p-value greater than 0.05 as well as *pCVB-eGFP* were filtered out. The *ggvenn* function in the ggplot2 package was used to create venn diagrams and identify unique as well as shared genes for each pairwise comparison. The *EnhancedVolcano* function was used to generate volcano plots for each pairwise comparison.

The progeny package was leveraged to compare signaling pathway activity across conditions within each timepoint.^29^ In summary, the *progeny* function was used to compute pathway activity scores from the corresponding Seurat object. *ScaleData* was applied to scale and center the PROGENy scores. The scaled scores were then transformed into a data frame and matched with umi IDs and conditions. A summary table depicting the pathway names, conditions, and mean as well as standard deviations for the progeny activity scores, which was calculated with the *summarise* function from the plyr package, was created. *pheatmap* was then used on the summary table to graph the mean PROGENy activity scores for each pathway.

#### Analyses of CVB3 infected single cell populations

A violin plot depicting *pCVB-eGFP* transcript presence in each cell type was generated via *VlnPlot*.

To calculate CVB3 infection rates, *subset* was used to isolate the CVB3 and Control conditions from the integrated dataset. GFP positive cells were then identified within the CVB3 conditions by using the *WhichCells* function to generate a list of cells with a *pCVB-eGFP* expression level greater than 0. The list was utilized to categorize cells within the isolated conditions into GFP Positive, GFP Negative, and Control cells using a *while* loop. The resulting vector was added to the subsetted Seurat object. New Seurat objects of data isolated from either the 24 hr CVB3 or 48 hr CVB3 conditions were generated. The timepoint datasets were then furthered subsetted into GFP Positive or GFP Negative groups. For each timepoint, the *table* function was finally used to get the number of GFP positive or negative cells across the various cell populations. The infection rate for each cell population was calculated by dividing the number of GFP positive cells by the corresponding total cell number.

To determine the number of differentially expressed genes between CVB3 and Control conditions for each cell population, *subset* was used to isolate each cell population from the integrated dataset. For each time point within these Seurat objects, CVB3 and Control conditions were compared using the *FindMarkers* with logfc.threshhold set to 0 to identify differentially expressed genes. Genes with an adjusted p-value greater than 0.05 were filtered out. Excel csv files were then generated from the corresponding gene lists and the number of rows within these files was used to graph the number of differentially expressed genes for each cell population at the corresponding timepoint.

The lists of differentially expressed genes generated when comparing CVB3 to Control conditions for each cell population were then leveraged to conduct gene ontology analyses using Enrichr on either upregulated or downregulated genes.^101–103^ Gene ontology themes were then generated based on the general categories (i.e. Cell structures/organelles, signaling pathways, cell processes) statistically significant gene ontology terms (adjusted p-value < 0.05) represented. The various gene ontology terms associated with each theme were compiled alongside their associated genes, source cell type, and timepoint. A list of unique genes was generated for each theme to determine the corresponding number of associated differentially expressed genes. To generate heatmaps depicting cell-type specific gene expression for selected themes, a *for* loop was first run to subset each cell type, perform pairwise comparison for each cell type between CVB3 and Control conditions for the corresponding timepoint, and extract statistically significant (adjusted p-value < 0.05) log_2_ fold change values for genes from the corresponding unique gene list. The *pheatmap* function was then used to plot these fold changes.

#### Cell re-clustering analysis

Cell populations (α, β, and ductal) were isolated from the integrated dataset using the *subset* function. The datasets were re-clustered by UMAP embedding and using *FindNeighbors* and *FindClusters* to find the nearest cell neighbors based on gene expression. UMAPs were generated via the *DimPlot* function.

The CVB3 and Control conditions were subsetted from the corresponding cell-specific re-clustered integrated object. The RNA assay for each was accessed to help calculate the *pCVB-eGFP* expression value corresponding to the 50^th^ percentile using the *quantile* function. The CVB datasets for each cell type were then divided into bystander (*pCVB-eGFP* expression level = 0), low expression (*pCVB-eGFP* expression level less than 50^th^ percentile but greater than 0), and high expression (*pCVB-eGFP* expression level greater than or equal to the 50^th^ percentile) groups using *WhichCells*. The umi IDs within these three groups were leveraged to designate the cells in the corresponding isolated Seurat object containing the isolated CVB3 and Control datasets. More specifically, a *while* loop was used to designate the cells as either Low GFP, High GFP, or Control and store this information into a vector. The resulting vector was added to the corresponding Seurat object via *append*.

For Pearson correlation analysis, the RNA assay for each cell-specific re-clustered object was accessed to identify any outlier genes on a mean variability plot using *FindVariableFeatures. AverageExpression* was then used on the top 19,172 variable genes. Pearson correlation was run on these average activity scores using the *cor* function and results were plotted using *heatmap*.*2*.

A list of differentially expressed genes when comparing Low GFP, High GFP, and Control cells was generated with *FindAllMarkers*. After removing *pCVB-eGFP* from each list, *AverageExpression* was used on the top 20 upregulated genes for each group. The results were plotted using *heatmap*.*2*.

#### Pseudotime trajectory analysis

Pseudotime analysis was performed using Monocle 3 version 1.2.9.^31–34^ The Seurat object containing Low GFP, High GFP, and Control groups from each cell type was converted to CellDataSet monocle-compatible file. Pseudotime trajectory was determined using *cluster_cells, learn_graph*, and *order_cells*. The trajectory root in *order_cells* was chosen objectively by *get_earliest_principal_node*, which picks a root by identifying a node heavily occupied by Control cells. The pseudotime values were then extracted and added to the corresponding Seurat object. *FeaturePlot* was used to generate pseudotime trajectory UMAPs. Genes dynamically changing as a function of pseudotime were found through *graph_test* with the *neighbor_graph* argument set to “principal_graph” and by excluding features with a low Moran’s *I* scores (I > 0.15). The genes and their dynamic expression over pseudotime were graphed using *Heatmap* with *K* means parameter set to two groups.

### Electron microscopy

Whole islets were fixed with 2.5% formaldehyde, 2.5% glutaraldehyde in sodium cacodylate buffer at pH 7.4 overnight at 4 °C and shipped to Harvard Medical School Electron Microscopy Core Facility for processing and sectioning. The sectioned samples were imaged with a JEOL 1200EX Microscope at 80 kV. Mitochondrial size was quantified using ImageJ.

### Reactive oxygen species detection

Reactive oxygen species levels were measured using a dihydroethidium assay kit (Abcam, ab236206). Briefly, treated whole islets were single cell dispersed in warm TrypLE at 37 °C for 15 min. The cells were transferred to a V-bottom 96-well plate (Corning, 3894) at a density of 80,000 cells/well and centrifuged at 400 x *g* for 1 min. After careful aspiration, 150 μL of Cell-Based Assay Buffer were added to each well and the cells were centrifuged as previously mentioned. We added 130 μL of ROS Staining Buffer to each well and 10 μL of N-acetyl Cysteine solution to designated negative controls. The plate was covered and incubated for 30 min at 37 °C protected from light. We added 10 μL of antimycin A solution were to designated positive controls, which was followed by an additional hour of incubation at 37 °C protected from light. The plate was centrifuged as previously mentioned, and the media was aspirated without disrupting the pellet. The pellet was resuspended in 100 μL of Cell-Based Assay Buffer and transferred to a black, clear bottom 96-well plate (Costar, 3875). Fluorescence was measured using a Synergy H1 microplate reader (BioTek) set to an excitation wavelength of 500 nm and an emission wavelength of 585 nm.

### Mitochondrial respiration and bioenergetics

Seahorse XF sensor cartridges (Agilent) were hydrated with sterile water at 37 °C without CO_2_ 24 hr before the run. On the day of the run, the injection ports were submerged in XF calibrant solution (Agilent, 100840-000) for at least 45 min. About 20-30 islets were collected for each technical replicate, washed with 1X PBS (Corning, 21-030-CV), and plated into a well of Seahorse XF24 Islet Capture Microplates (Agilent, 101122-100) with capture screens. Each well contained Seahorse XF RPMI media at pH 7.4 (Agilent, 103576-100) and 20 mM glucose. The number of islets per well was determined by counting under a microscope. The islets were then equilibrated at 37 °C without CO_2_ for at least 1 hr. The cells and sensors were placed in a Seahorse XFe24 extracellular flux analyzer (Agilent). After 5 baseline measurements, the cells were treated with sequential injections of 3 μM oligomycin (Sigma, O4876), 1 μM carbonyl cyanide 4-(trifluoromethoxy) phenylhydrazone (FCCP), and 1 μM rotenone (Sigma, R8875) together with 2 μM antimycin A (Sigma, A8674).

The resulting data was normalized to islet number and analyzed following steps and code adjusted from Yépez et al. (2018).^30^ Briefly, the combined patient dataset for each timepoint was uploaded into R using *fread. add_outlier_col* was utilized to identify and remove well outliers first followed by individual measurement outliers. The bioenergetic values were then measured via *compute_bioenergetics* using the LR_ao method. A comparison table is generated with *create_comp_table* to compare each treatment group to the control in downstream analyses. Finally, *stat_test_OCR* is run on the bioenergetics values to perform statistical testing between samples.

### HUES8 culture

HUES8 human embryonic stem cells (hESCs) were cultured mTeSR1 (Stem Cell Technologies, 05850) media and incubated at 37 °C with 5% CO_2_. The cells were passaged every 3-4 days by washing with 1X PBS and incubating with TrypLE Express at 0.2 mL/cm^2^ for 7 min or less at 37 °C. An equal volume of mTeSR1 supplemented with 10 μM Y-27632 (Pepro Tech, 129382310MG) was then added. Dispersed cells were counted with a Vi-Cell XR (Beckman Coulter) and centrifuged at 300 x *g* for 3 min. The supernatant was aspirated, and cells were resuspended in supplemented mTSeR1 before plating onto Matrigel-coated plates at a cell density of 0.8 × 10^5^ cells/cm^2^ for propagation. The cells were cultured in mTSeR1 with Y-27632 for 24 hr. The cells were then fed daily with mTSeR1 without Y-27632.

### HUES8 Differentiation

HUES8 hESCs were differentiated following our previously published protocol.^36,37^ Briefly, hESCs were single cell dispersed and seeded at 0.63 × 10^6^ cells/cm^2^ with mTSeR1 supplemented with Y-276352. After 24 hr, the media was replaced with differentiation base media with the corresponding factors. On the 7^th^ day of stage 6, the cells are single cell dispersed and seeded at 5 × 10^6^ cells/well of a 6-well plate with 5 mL of the corresponding media. The cells are then placed on an Orbi-Shaker at 100 RPM. Stem cell-derived islets were used for assays between day 15-20 in stage 6.

### Lentivirus production and transduction

Plasmids containing the short hairpin RNA (shRNA) sequences of interest were ordered as glycerol stocks and then grown. Plasmid DNA was isolated using the QIAprep Miniprep kit (Qiagen, 27115) and then transformed into One Shot™ Stbl3™ Chemically Competent *Escherichia coli* cells (Invitrogen, C737303). Single colonies were picked after 18 hr, cultured, and subjugated to the Qiagen Maxi plus kit (Qiagen, 12981) for DNA extraction. Lentiviral particles were generated using the Lenti-X 293T cells (Takara, 632180) cultured in 10 cm plates (Falcon, 353003) with Dulbecco’s modified Eagle medium, 10% heat-inactivated feral bovine serum (MilliporeSigma, F4135), and 0.01 mM sodium pyruvate (Corning, 25-000-CL). Once confluent, the Lenti-X 293T cells were transfected with 6 μg of plasmid DNA, 4.5 μg of psPAX2 (Addgene, 12260), 1.5 μg pMD2.G (Addgene, 12259), and 48 μL of Polyethylenimine Max (Polysciences, 24765-2). Media was changed after 16 hr. Virus containing supernatant was collected at 96 hr post-transfection, concentrated using the Lenti-X concentrator (Takara, 631232), and tittered using Lenti-X GoStix Plus (Takara, 631280). SC-islets were transduced with lentiviral vectors containing shRNAs targeting either *GFP* (Control) or *MIR7-3HG* on Stage 6 Day 7 during aggregation at the corresponding MOI. After 24 hr, the cells were washed with media to remove excess viral particles.

### Viability assays

Cell viability was measured by multiplexing the CellTiter-Fluor Cell Viability Assay (Promega, G6080) with the Caspase-Glo 3/7 Assay System (Promega, G8090). Briefly, stem cell-derived islets were single cell dispersed and plated into 96-well white with clear bottom plates (Corning, 3903) at a cell density of 4 × 10^4^ cells/well. After 24 hr, the cells treated with either endotoxin-free water, or CVB3-eGFP (titer = 1.63 × 10^8^ pfu/mL) at a MOI of 0.1. The cells were washed with 1X PBS 48 hr post infection and 80 μL of culture media were added per well. Each well then received 20 μL of the viability assay reagent and the plate was gently swirled for mixing. The cells were incubated at 37 °C for 30 min prior to measuring fluorescence using a Synergy H1 microplate reader (BioTek). Afterwards, 100 μL of the Caspase-Glo 3/7 reagent were added followed by a 30 min incubation at room temperature in the dark. The Synergy H1 microplate reader was used to measure luminescence.

### Statistical Analysis

The sample sizes for scRNA-sequencing contain 4 separate patients, 3 of which were independently processed for hashing using the TotalSeq™ antibody cocktails. Statistical analyses were calculated using GraphPad Prism and R. GraphPad Prism was also utilized to identify any outliers using the ROUT method. Wilcoxon Rank Sum test in conjunction with Bonferroni correction were used to obtain statistically significant DEGs with an adjusted p-value < 0.05. We used Ordinary One-way ANOVA with Tukey’s multiple comparisons for rt-qPCR, mitochondrial area, and ROS assays. P-values for mitochondrial OCR were obtained using a Student’s t-test as detailed by Yépez et. al.^30^ TEM images were obtained using human cadaveric islets from two patients. ROS measurements were made using islets from one patient. Islets from the 3 hashed patients were also used for mitochondrial respiration and bioenergetics assays. SC-islets from two independent differentiation batches were used for *shMIR7-3HG* #1. Three independent differentiation batches were used for all assays involving *shMIR7-3HG* #2. P-values are marked as follows: non-significant (ns) > 0.05, * < 0.05, ** < 0.01, *** < 0.001, and **** < 0.0001.

## Extended Data Figures

**Extended Data Fig. 1.**
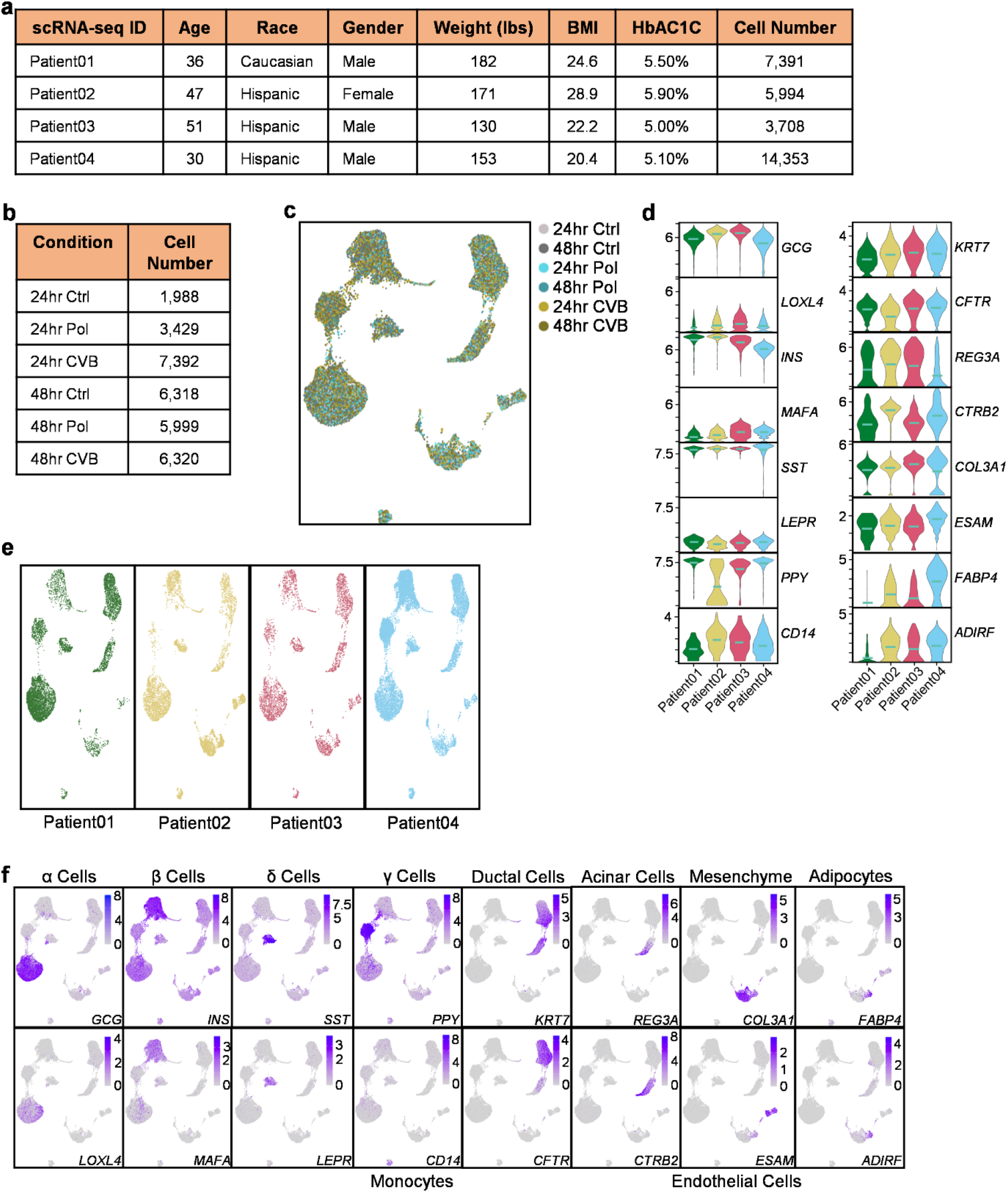
Single-cell RNA-seq similarly captures all the major pancreatic cell types across all patients. **a**, Patient information from scRNA-seq replicates. **b**, Number of total cells found in each treatment condition after quality check of scRNA-seq dataset. **c**, UMAP depicting distribution of cells for each condition in the scRNA-seq dataset. **d**, Violin plots of cell marker expression across patients in control conditions for both timepoints. **e**, UMAPs showing cluster distribution across patients. **f**, Feature plots depicting cell marker expression within clusters.

**Extended Data Fig. 2.**
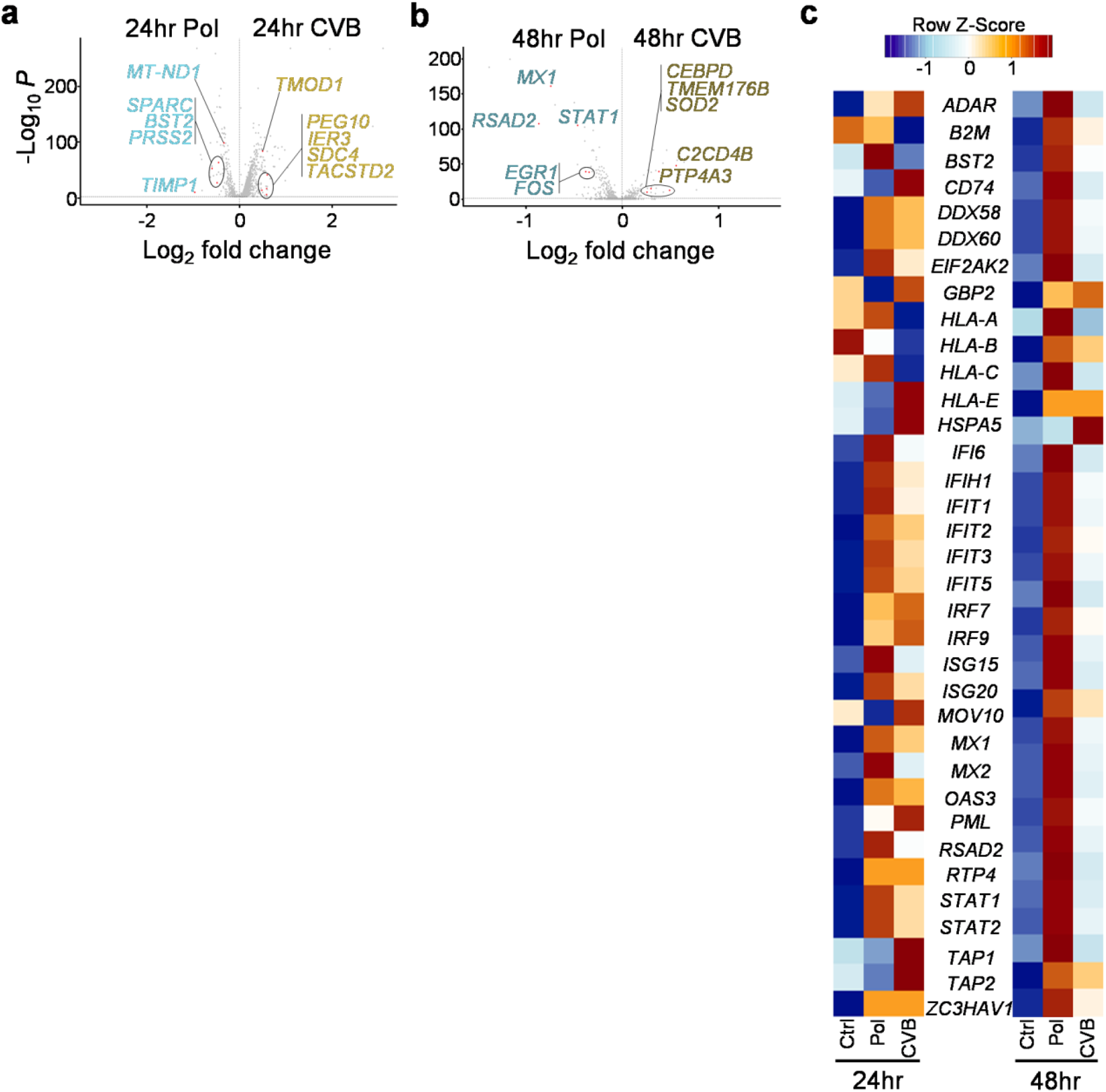
Single-cell RNA-seq confirms CVB3 infection and reveals major transcriptional differences between CVB3 and poly(I:C) treated islets. **a**, Volcano plot highlighting the differences in gene expression when comparing poly(I:C) treated to CVB3 infected islets at 24 hr (total variables = 2753). **b**, Volcano plot highlighting the differences in gene expression when comparing poly(I:C) treated to CVB3 infected islets at 48 hr (total variables = 438). **c**, Heatmap of genes associated with type 1 IFN signaling in primary human islets highlighting temporal differences between poly(I:C) and CVB3.

**Extended Data Fig. 3.**
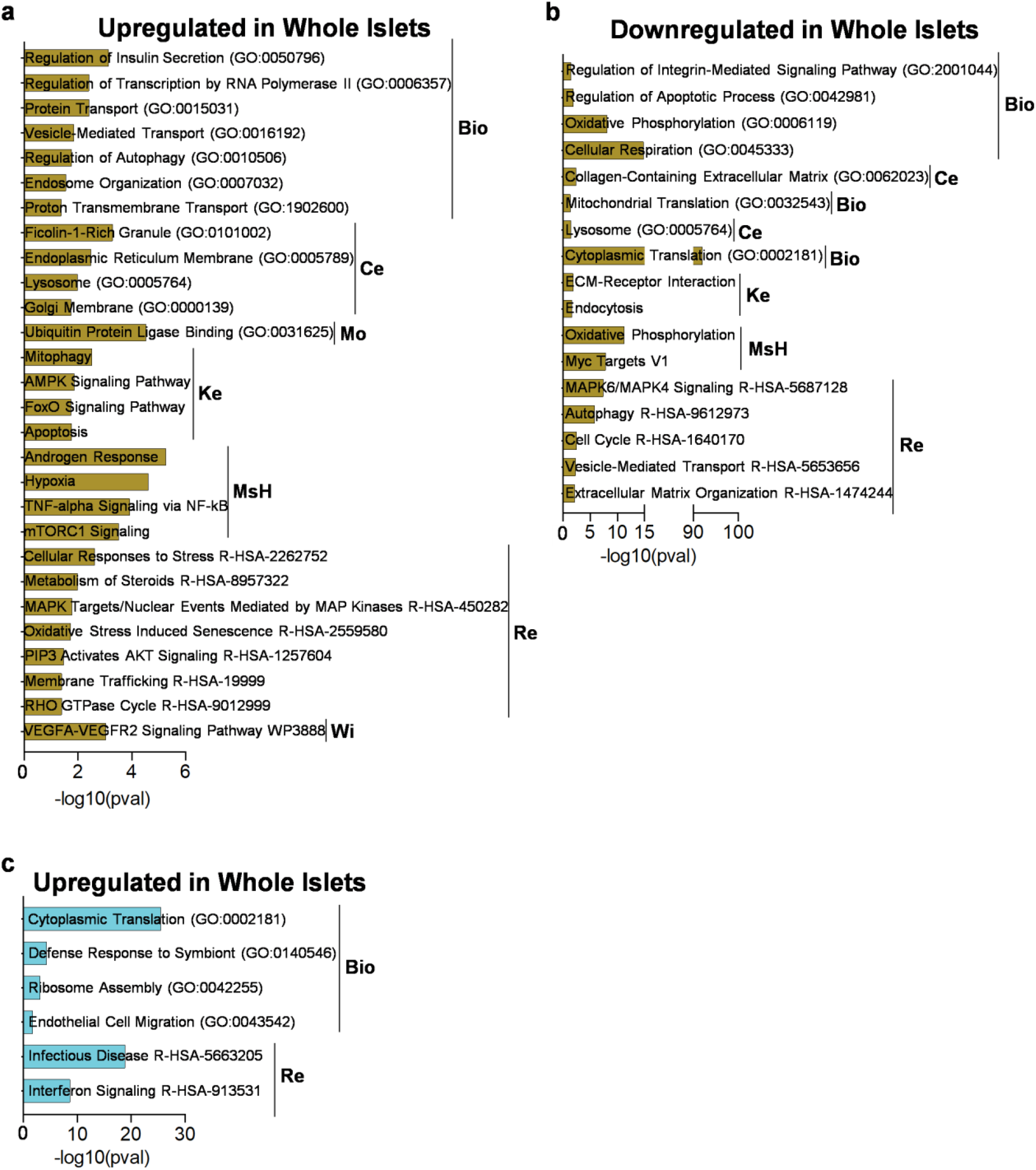
Gene ontology (GO) analyses of whole islets comparing control conditions to either CVB3 or poly(I:C) show poly(I:C) induces a less complex transcriptional response at 24 hr. **a**, GO analysis of pathways enriched in genes upregulated in CVB compared to control after 24 hr (Ke = KEGG 2021, Mo = Molecular Function 2023, Ce = Cellular Component 2023, Wi = WikiPathway 2021, Bio = Biological Process 2023, MsH = MSigDB Hallmark 2020, Re = Reactome 2022). **b**, GO analysis of pathways enriched in genes downregulated in CVB3 compared to control after 24 hr. **c**, GO analysis of pathways enriched in genes upregulated in poly(I:C) compared to control after 24 hr.

**Extended Data Fig. 4.**
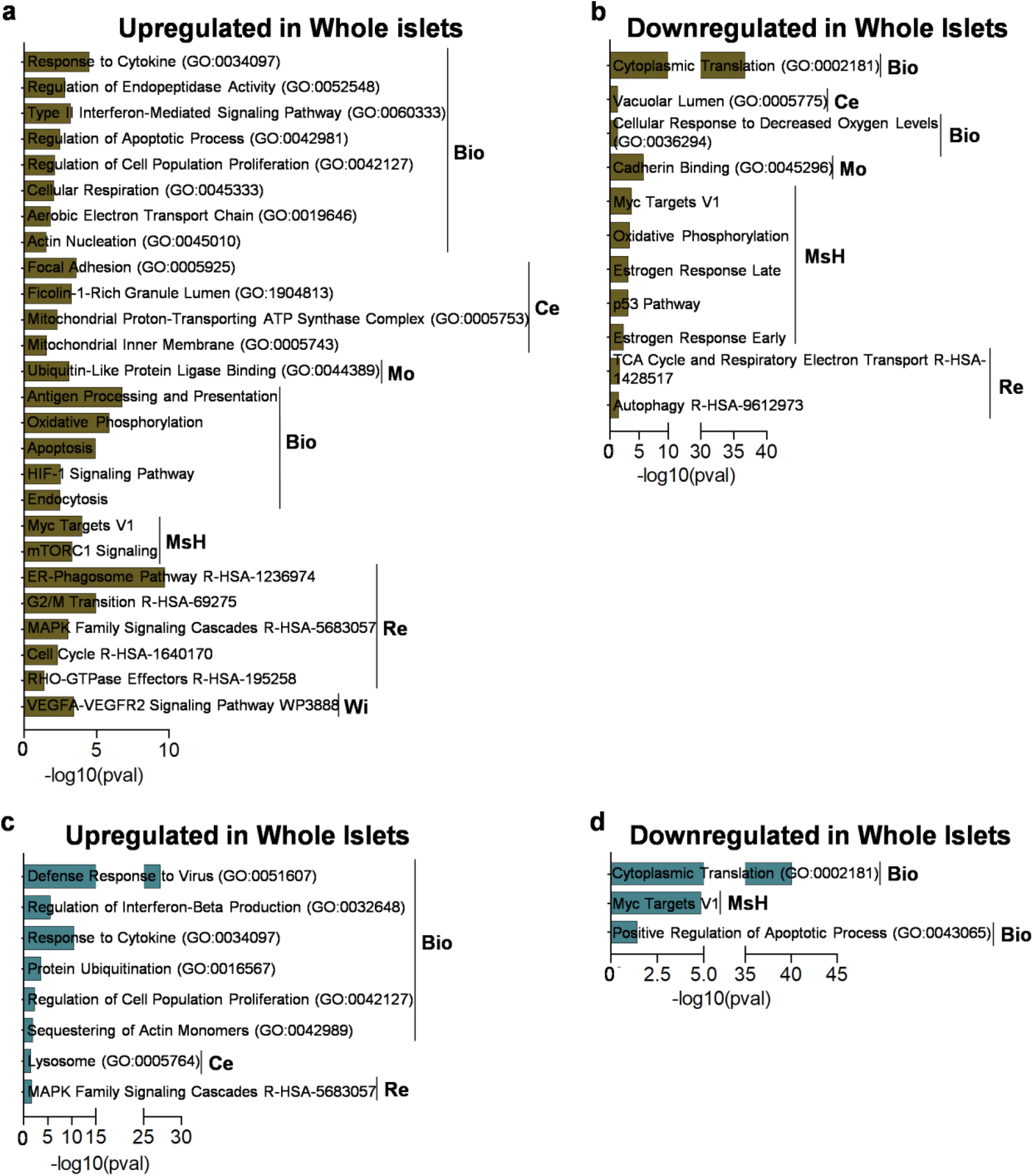
GO analyses of whole islets comparing control conditions to either CVB3 or poly(I:C) show poly(I:C) induces a less complex transcriptional response at 48 hr. **a**, GO analysis of pathways enriched in genes upregulated in CVB3 compared to control after 48 hr (Ke = KEGG 2021, Mo = Molecular Function 2023, Ce = Cellular Component 2023, Wi = WikiPathway 2021, Bio = Biological Process 2023, MsH = MSigDB Hallmark 2020, Re = Reactome 2022). **b**, GO analysis of pathways enriched in genes downregulated in CVB3 compared to control after 48 hr. **c**, GO analysis of pathways enriched in genes upregulated in poly(I:C) compared to control after 48 hr. **d**, GO analysis of pathways enriched in genes downregulated in poly(I:C) compared to control after 48 hr.

**Extended Data Fig. 5.**
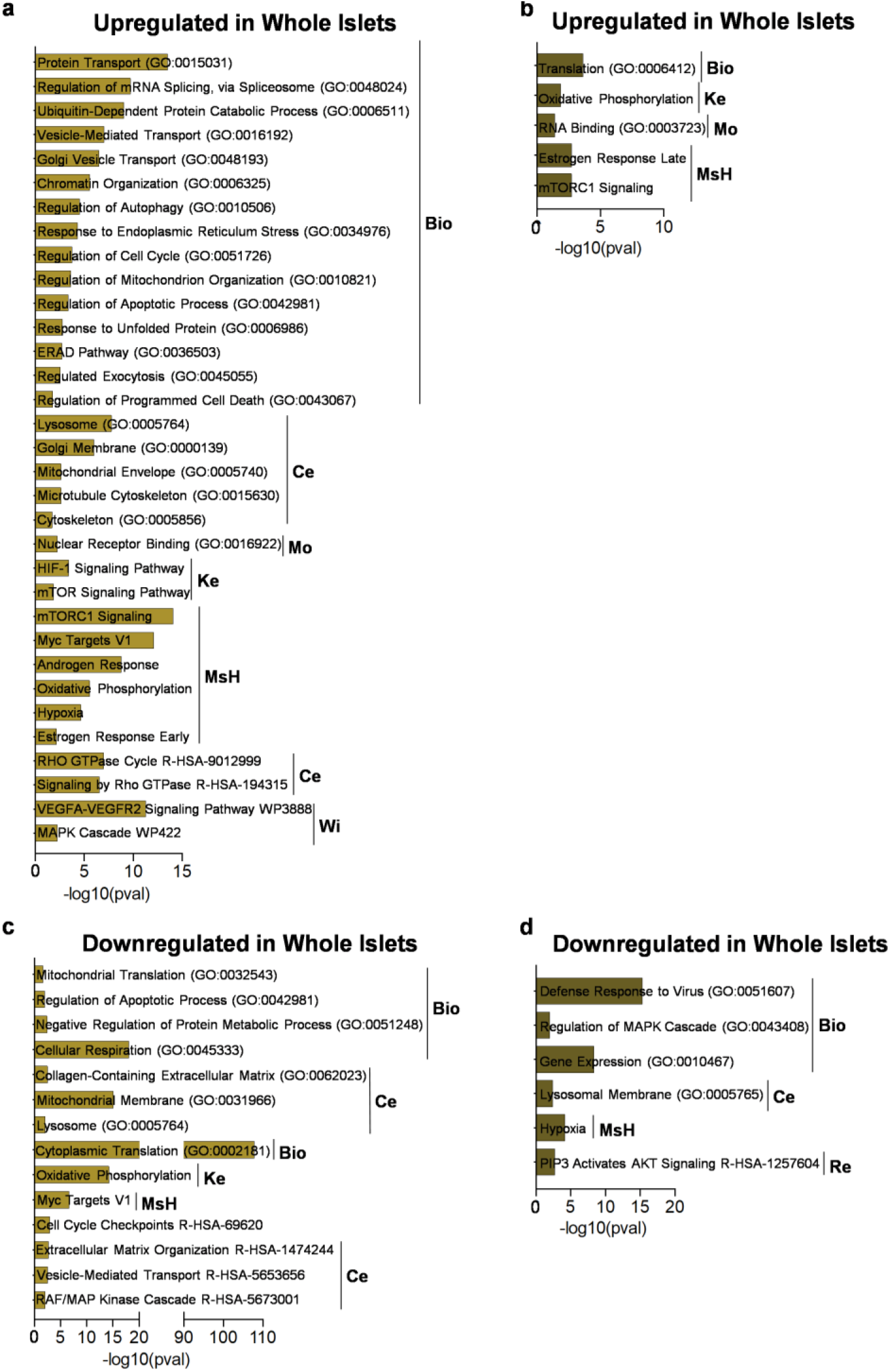
GO analyses of whole islets comparing CVB3 conditions to poly(I:C) show poly(I:C) induces a less complex transcriptional response. **a**, GO analysis of pathways enriched in genes upregulated in CVB3 compared to poly(I:C) after 24 hr (Ke = KEGG 2021, Mo = Molecular Function 2023, Ce = Cellular Component 2023, Wi = WikiPathway 2021, Bio = Biological Process 2023, MsH = MSigDB Hallmark 2020, Re = Reactome 2022). **b**, GO analysis of pathways enriched in genes upregulated in CVB3 compared to poly(I:C) after 48 hr. **c**, GO analysis of pathways enriched in genes downregulated in CVB3 compared to poly(I:C) after 24 hr. **d**, GO analysis of pathways enriched in genes downregulated in CVB3 compared to poly(I:C) after 48 hr.

**Extended Data Fig. 6.**
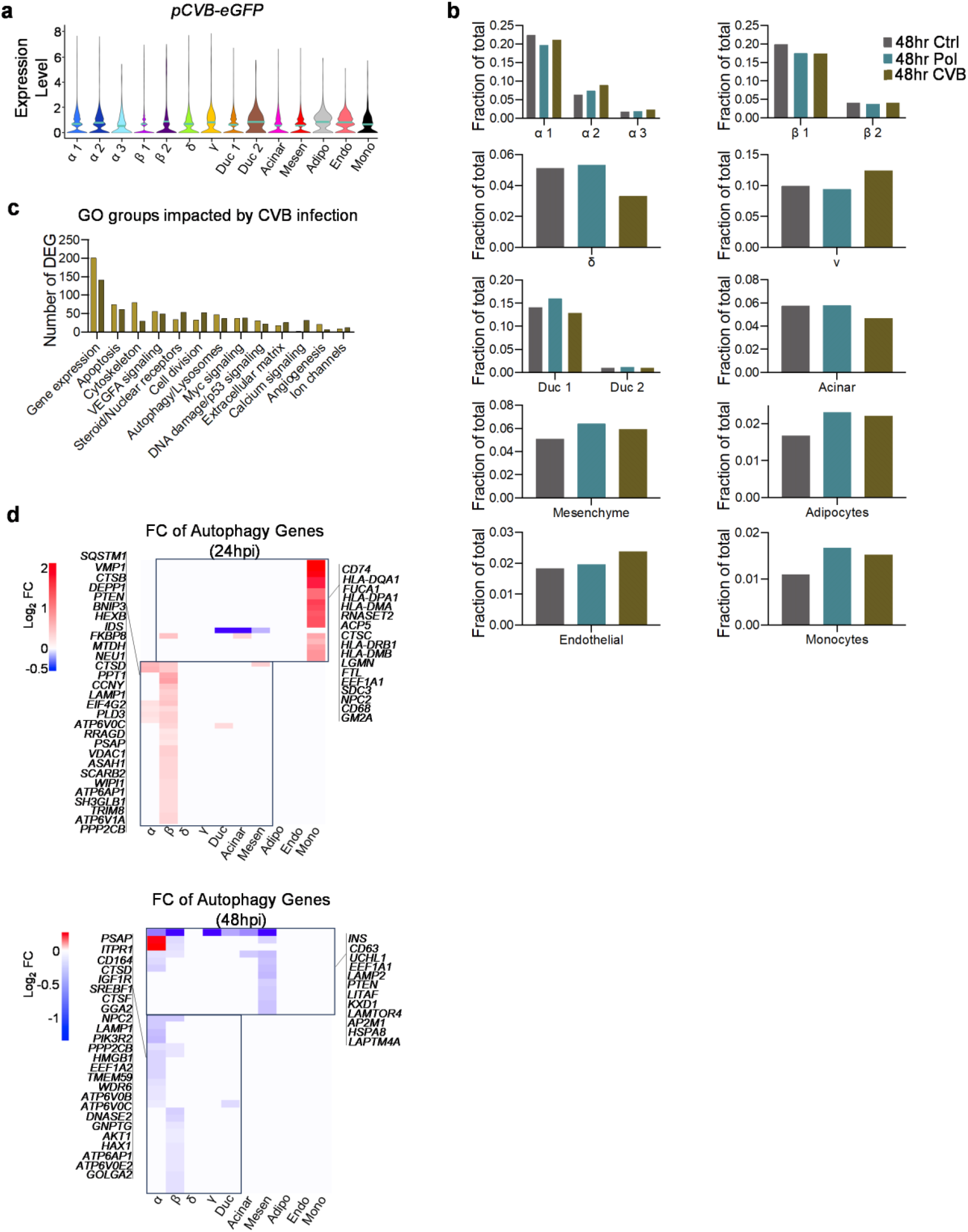
scRNA-seq reveals unique cell-type specific responses to CVB3 infection despite similar infection rates. **a**, Violin plot of CVB3-eGFP expression across all cell types. **b**, Bar graphs of cell proportions in 48 hr conditions. **c**, Bar graph showing the number of differentially expressed genes for selected themes of common gene ontology terms for each CVB3 timepoint compared to corresponding control. **d**, Heatmaps depicting log_2_ fold change of autophagy-associated genes when comparing 24 hr CVB3 vs control and 48 hr CVB3 vs control.

**Extended Data Fig. 7.**
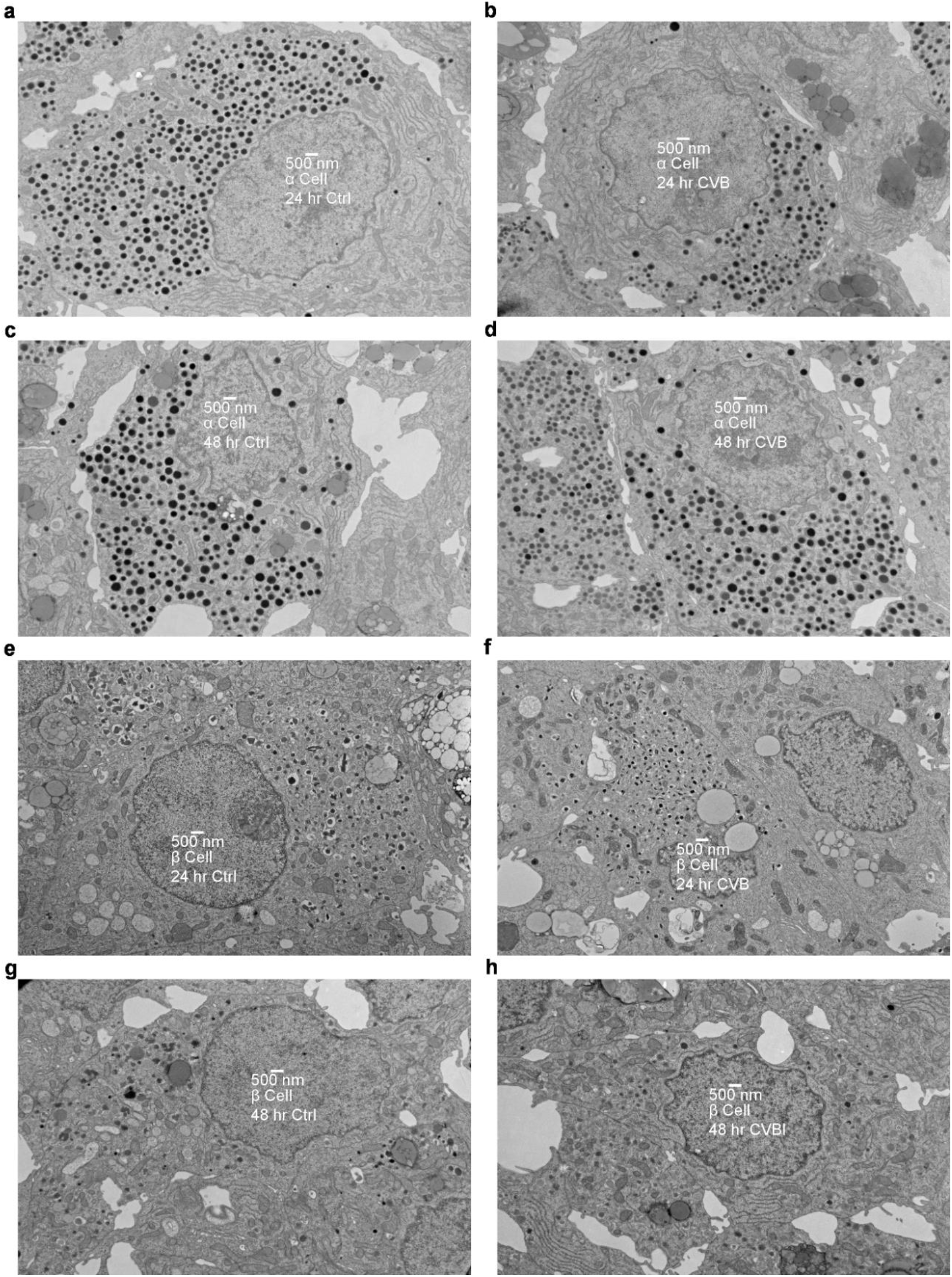
TEM images of α and β cells taken at 4000x direct magnification. **a**, TEM image of a control α cell at 24 hr. **b**, TEM image of a CVB3 infected α cell at 24 hr. **c**, TEM image of a control α cell at 48 hr. **d**, TEM image of a CVB3 infected α cell at 48 hr. **e**, TEM image of a control β cell at 24 hr. **f**, TEM image of a CVB3 infected β cell at 24 hr. **g**, TEM image of a control β cell at 48 hr. **h**, TEM image of a CVB3 infected β cell at 48 hr.

**Extended Data Fig. 8.**
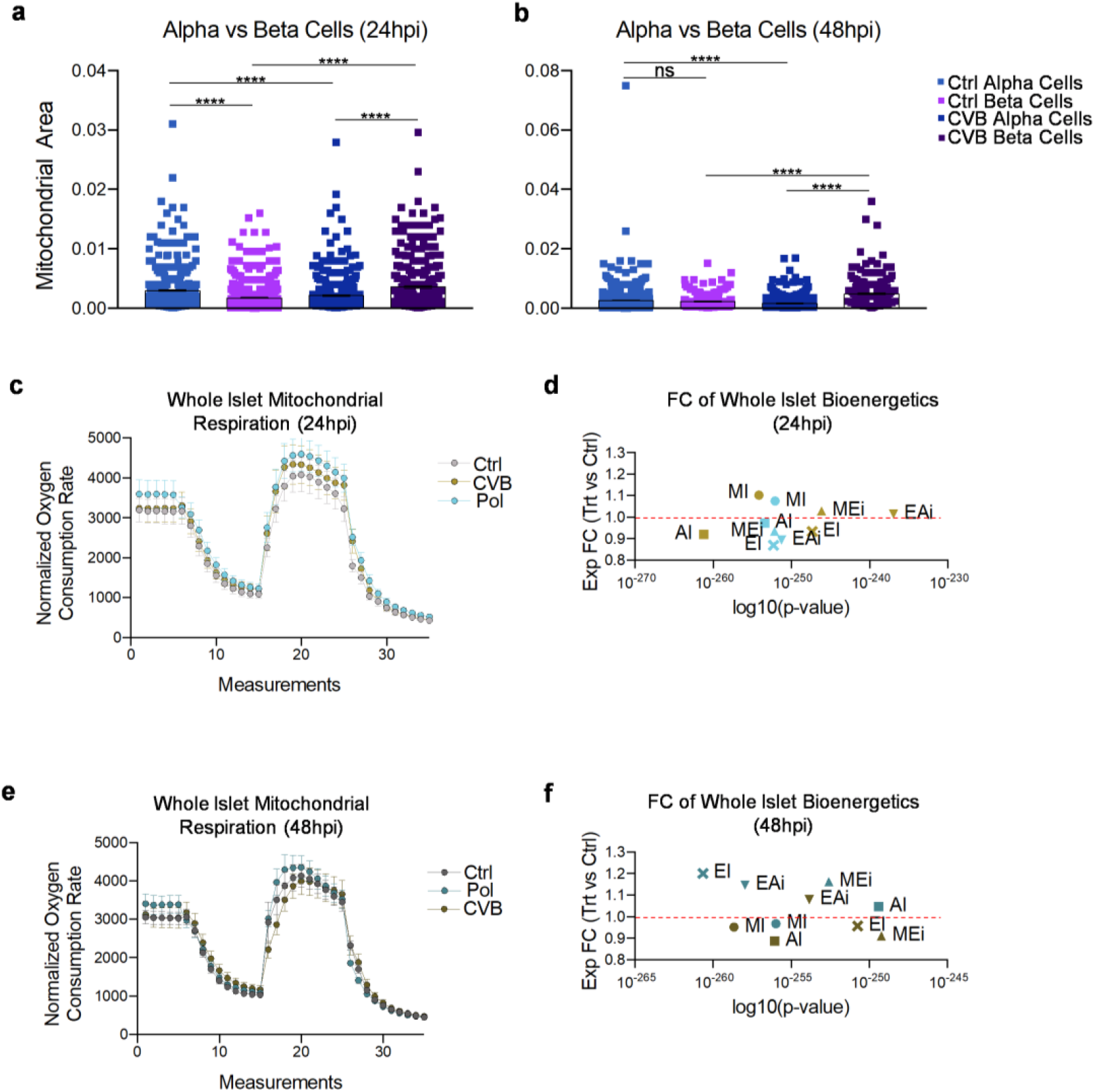
CVB3 infection induces mitochondrial dysfunction in primary human islets at 48 hr. **a**, Mitochondrial area of α cells vs β cells at 24 hr based off measurements made from TEM images using Fiji. (24 hr α Ctrl, n = 494; 24 hr α CVB, n = 632; 24 hr β Ctrl, n = 669; 24 hr β CVB, n = 476). ** = One-way ANOVA with Tukey’s multiple comparisons. **b**, Mitochondrial area of α cells vs β cells at 48 hr based off measurements made from TEM images using Fiji. (48 hr α Ctrl, n = 668; 48 hr α CVB, n = 815; 48 hr β Ctrl, n = 357; 48 hr β CVB, n = 210). ** = One-way ANOVA with Tukey’s multiple comparisons. **c**, Seahorse XF test measuring mitochondrial respiration at 24 hr of primary human islets from patients 01, 02, and 03 before and after injection with Oligomycin, FCCP, and Antimycin A/rotenone (Data normalized to islet number per well). **d**, Exponential fold change (FC) of bioenergetics comparing poly(I:C) and CVB3 to control at 24 hr. (EI = ETC-dependent OCR proportion, AI = ATPase-dependent OCR proportion, EAi = ETC-dependent proportion of ATPase-independent OCR, MI = Maximal over initial OCR FC, MEi = Maximal over ETC-independent OCR FC). ** (Student’s t-test). **e**, Seahorse XF test measuring mitochondrial respiration of primary human islets from patients 01, 02, and 03 before and after injection with Oligomycin, FCCP, and Antimycin A/rotenone (Data normalized to islet number per well). **f**, Exponential FC of bioenergetics comparing poly(I:C) and CVB3 to control at 48 hr. ** (Student’s t-test).

**Extended Data Fig. 9.**
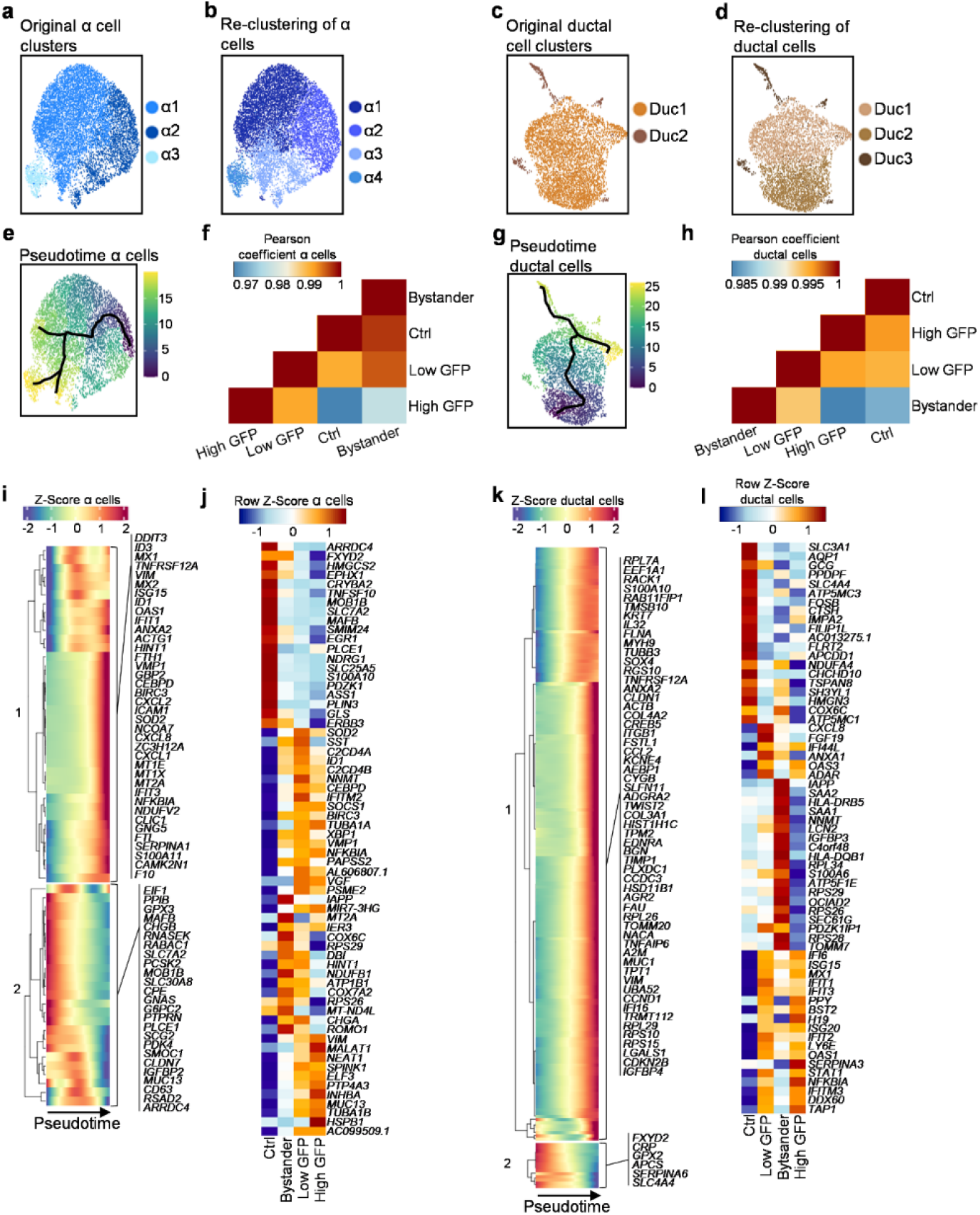
α and ductal cells exhibit differential transcriptional responses to CVB3 infection dependent on viral load. **a**, UMAP showing original distribution of α cell populations (subset of α cell populations from all conditions). **b**, UMAP showing subpopulations upon re-clustering of α cell populations only. **c**, UMAP showing original distribution of ductal cell populations (subset of ductal cell populations from all conditions). **d**, UMAP showing subpopulations upon re-clustering of ductal cell populations only (subset of ductal cell populations from all conditions). **e**, UMAP showing the pseudotime trajectory of α cell subpopulations (subset of α cell populations from control and CVB3 conditions). **f**, Pearson correlation plot of α cell subpopulations grouped by viral load (subset of α cell populations from control and CVB3 conditions). **g**, UMAP showing the pseudotime trajectory of ductal cell subpopulations (subset of ductal cell populations from control and CVB3 conditions). **h**, Pearson correlation plot of ductal cell subpopulations grouped by viral load (subset of ductal cell populations from control and CVB3 conditions). **i**, Pseudotime trajectory heatmap showing the dynamic changes of α cell gene expression upon CVB3 infection. **j**, Heatmap of the top upregulated genes in control, bystander, low GFP, and high GFP α cells. **k**, Pseudotime trajectory heatmap showing the dynamic changes of ductal cell gene expression upon CVB3 infection. **i**, Heatmap of the top upregulated genes in control, bystander, low GFP, and high GFP ductal cells.

**Extended Data Fig. 10.**
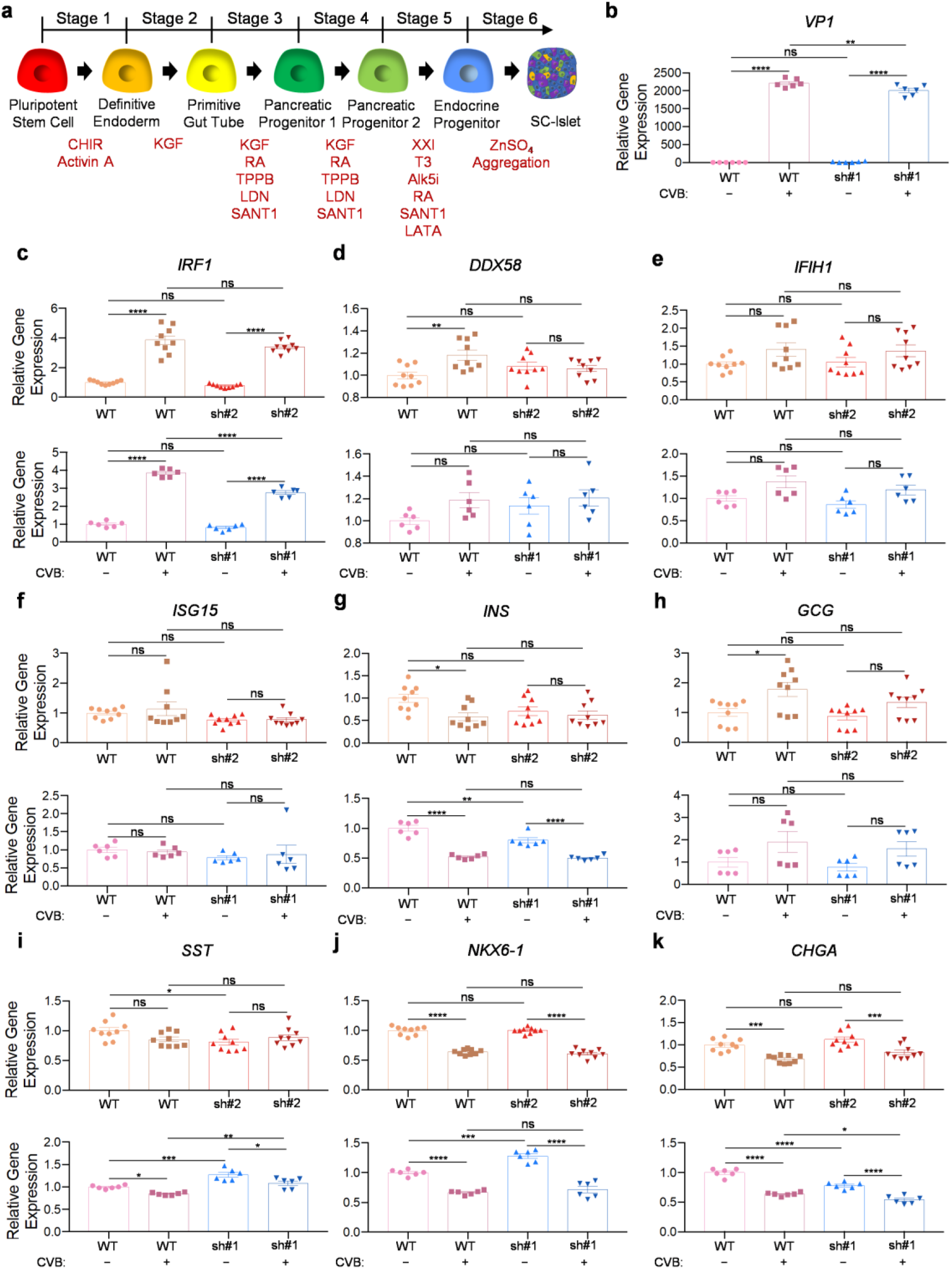
*MIR7-3HG* regulates apoptosis leading to lower viral load despite no major impact on type 1 IFN and identity gene expression in SC-islets infected with CVB3. **a**, Schematic of protocol adapted from Hogrebe et al. (2021) to differentiate hPSCs into SC-islets. **b**, rt-qPCR of viral antigen expression upon knockdown of *MIR7-3HG* and CVB infection of SC-islets. **c-f**, rt-qPCR of different type 1 IFN-associated genes upon *MIR7-3HG* knockdown with two shRNAs. **g-k**, rt-qPCR of different islet identity genes upon *MIR7-3HG* knockdown with two shRNAs. ** = two-way ANOVA with Tukey’s multiple comparisons.

## Notes

### Summary of Updates

Figures 1 and 2 were inadvertently switched in the original version. The revision has corrected this error.

